# A common computational and neural anomaly across mouse models of autism

**DOI:** 10.1101/2024.05.08.593232

**Authors:** Jean-Paul Noel, Edoardo Balzani, Luigi Acerbi, Julius Benson, The International Brain Laboratory, Cristina Savin, Dora E. Angelaki

## Abstract

Computational psychiatry has suggested that humans within the autism spectrum disorder (ASD) inflexibly update their expectations (i.e., Bayesian priors). Here, we leveraged high-yield rodent psychophysics (n = 75 mice), extensive behavioral modeling (including principled and heuristics), and (near) brain-wide single cell extracellular recordings (over 53k units in 150 brain areas) to ask (1) whether mice with different genetic perturbations associated with ASD show this same computational anomaly, and if so, (2) what neurophysiological features are shared across genotypes in subserving this deficit. We demonstrate that mice harboring mutations in *Fmr1*, *Cntnap2*, and *Shank3B* show a blunted update of priors during decision-making. Neurally, the differentiating factor between animals flexibly and inflexibly updating their priors was a shift in the weighting of prior encoding from sensory to frontal cortices. Further, in mouse models of ASD frontal areas showed a preponderance of units coding for deviations from the animals’ long-run prior, and sensory responses did not differentiate between expected and unexpected observations. These findings demonstrate that distinct genetic instantiations of ASD may yield common neurophysiological and behavioral phenotypes.

## Main

Autism spectrum disorder (ASD) is a neurodevelopmental condition of unknown etiology. The disorder is characterized by abundant heterogeneity, both at biological (e.g., genetics and neurochemistry^1–4^) and behavioral scales (e.g., anomalies across social, communicative, perceptual, and motor domains^5–7^). This diversity hinders our ability to understand and ultimately treat the condition.

To address the phenotypic heterogeneity, computational psychiatry^8–17^ has recently cast ASD with the language of probabilistic inference^18^ and suggested that individuals within the spectrum (1) have attenuated expectations^19^ (2) inflexibly weight predictions relative to sensory observations^20^ and/or (3) over-estimate the volatility of their sensory environment and thus are less surprised by statistically unlikely events^21^. Critically, each of these computational accounts suggest that the learning of context-dependent statistical regularities (i.e., “contextual priors^22^”) is abnormal (e.g., slow^23, 24^) in ASD. This convergence on a putative computational deficit in humans with ASD supposes an opportunity to now account for biological heterogeneity.

We attempt to bridge the gap between the recent computational accounts of ASD and its neurobiological and genetic instantiation. Namely, we hypothesized that if a single computation (e.g., aberrant prior updating) may account for myriad of behavioral symptoms in ASD^19–24^, then the distinct genetic makeups of the condition may similarly (1) all express this computational deficit, and (2) share a common underlying neurophysiological profile subserving the computational deficit. We test this hypothesis across three different monogenetic mouse models of ASD (*Fmr1*^25^, *Cntnap2*^26^, and *Shank3B*^27^) by leveraging a standardized high-yield visual detection task requiring the use of priors^28^, behavioral modeling of online adaptation to changes in statistical regularities^29, 30^, and large-scale neurophysiological recordings with high-density silicon probes^31^.

The results demonstrate that all tested genetic mouse models of ASD underutilized adaptive statistical regularities during decision-making. These animals showed a blunted update of their priors^23, 24^ and were not surprised by statistically unlikely events^21^. Brain-wide extracellular recordings showed a neural convergence, wherein the relative weighting of prior encoding at the unit- and population-level shifted from sensory to frontal cortices in all mouse models of ASD. Likewise, we observed (1) a preponderance of units coding for deviations from the animals’ long-run prior (prior mean) and (2) a lack of sensory-driven prediction errors in frontal cortices of mouse models of ASD. Together, these results suggest that both global and local neural imbalances engender the inflexible updating of Bayesian priors in ASD. Globally, there is a shift in prior encoding from sensory to frontal areas. Locally, within frontal areas, there is a suppression of statistically surprising sensory observations and an outsized influence of signals coding for the prior mean. More broadly, the results demonstrate a common computational and neural anomaly across mouse models of ASD, suggesting that distinct genetic instantiations of the disorder may converge onto common behavioral and neurophysiological phenotypes.

## Results

### Reduced utilization of statistical regularities in mouse models of ASD

Wildtype mice (C57BL/6j, n = 15) and three genetic mouse models of ASD – *Fmr1*^-/y^ (n = 19), *Cntnap2^lacZ/+^*(n = 21), *Shank3B^+/-^* (n = 20) – were first trained on a prior-independent, two-alternative forced-choice visual detection task. We leveraged the standardized task and protocols from the International Brain Lab^28^ (**Fig. 1A**). Briefly, a grating of varying contrast was presented with equal probability to the left or right of a head-fixed mice. The animal then selected one of two choices (i.e., left or right) by turning a steering wheel coupled to the grating location. If the mouse brought the grating to the center of its visual field, it was rewarded with sucrose. Instead, if the animal moved the grating by an equivalent distance in the other direction (i.e., off-screen), it was penalized by a timeout (**Fig. 1B**, see *Methods* for detail).

**Figure 1.**
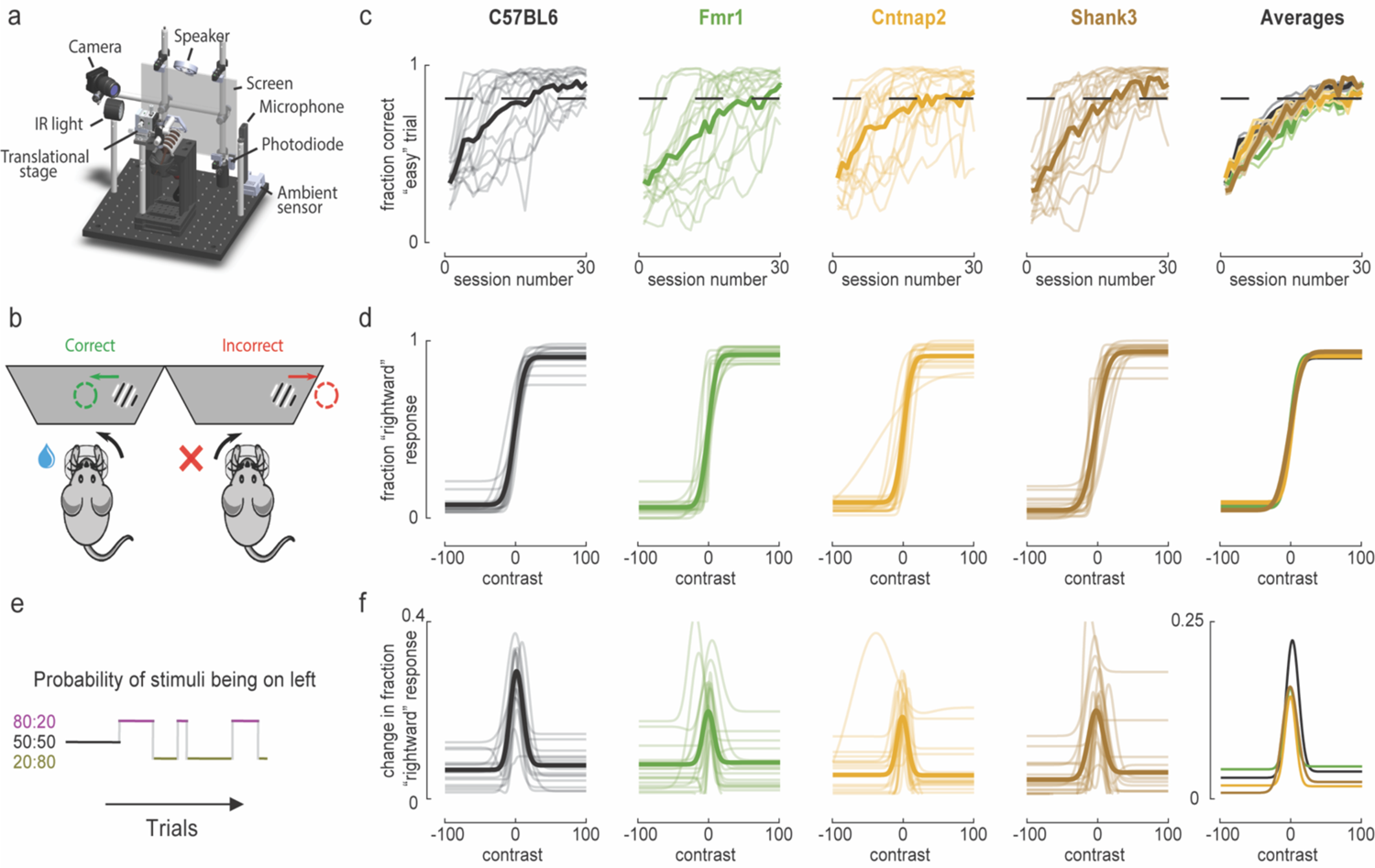
Reduced utilization of statistical regularities in mouse models of ASD. **A.** Rendering of the standardized behavioral apparatus allowing for the study of visually-guided decisions in rodents. **B.** Schematic of the task. **C.** Performance of control (C57BL6) and mouse models of ASD (*Fmr1*, *Cntnap2*, *Shank3*) on “easy” trials (i.e., 50% and 100% contrast) as a function of session number. Animals were considered trained when they performed at or above 80% (black dashed line) on “easy” trials (moving average over 3 sessions). **D**. Psychometric functions describing the fraction of rightward choices as a function of contrast and animal genotype. Negative contrasts denote stimuli on the left. Thin and transparent lines are individual animals, while thicker and opaque lines are averages. These psychometric curves are derived by combining all sessions after animals were considered proficient (average number of trials per animal = 1250.9). **E.** Schematic illustrating the “biased” version of the task. The fist 90 trials are “unbiased” in that gratings appear with equal probability on the left and right (50:50). Subsequently, blocks of varying length (range = 20-100 trials, decaying exponential such that the hazard rate was approximately constant) show gratings predominantly on the left or right (80:20 vs. 20:80; respectively in purple and gold). **F.** Change in the fraction of rightward responses as a function of block (rightward – leftward, fitted curves as in **d**), contrast, and animal genotype (average n trials per animal = 5645.0). Vertical axis (y) in the rightmost panel is compressed to show difference between wildtype animals (black) and mouse models of ASD (colored).

Mice of all genotypes learned this task (learned/total; *C57BL6*: 15/15; *Fmr1:* 17/19*; Cntnap2:* 19/21; *Shank3:* 18/20; χ^2^ test, p = 0.90). Further, provided that an animal became proficient at the task, it did so in an equal number of sessions (**Fig. 1C**, mean ± s.e.m.; *C57BL*: 12.9 ± 2.2; *Fmr1:* 16.8 ± 2.2*; Cntnap2:* 13.8 ± 2.4; *Shank3:* 12.8 ± 1.4) regardless of genotype (p = 0.52). At asymptotic behavior, psychometric performance on this prior-independent task was equal across genotypes (**Fig. 1D**, overall means ± s.e.m., bias: −1.97 ± 0.94; threshold: 14.6 ± 1.13; lapses: 0.06 ± 0.005; all p > 0.53), indicating that mouse models of ASD detect visual stimuli equally well to their control counterparts.

Following proficiency on the visual detection task, we introduced a dynamic prior. Sessions started with an unbiased block of trials (50:50 probability of stimuli being on the left or right) and then alternated between gratings being more frequent on the left or right visual field (80:20 vs. 20:80 probability; **Fig. 1E**). Importantly, the change in block (e.g., from leftward to rightward) was unsignaled and thus had to be inferred from the stream of stimulus statistics and sensory experience. Moreover, 0% contrast trials were rewarded according to the prior, and thus optimal performance on the task required that animals incorporate knowledge of the prior in making decisions.

To quantify the impact of this prior, we fit psychometric curves to responses during leftward and rightward biased blocks (**Eq. 1** in *Methods*, see **Fig. S1** for example animals). Then, we subtract these curves (fitted rightward – leftward psychometric curves; see^28^ for a similar approach). As expected, the prior was most informative when sensory evidence was weak, resulting in the largest difference between left- and right-prior blocks being at contrast zero (peak at 0.33% contrast, no difference across genotypes, p = 0.28; **Fig. 1F**). Most strikingly, the impact of the prior was greater in the control animals (differences in fraction “rightward” responses at contrast = 0, 0.27 ± 0.005) than the mouse models of ASD (**Fig. 1F**, *Fmr1:* 0.24 ± 0.004*; Cntnap2:* 0.20 ± 0.003; *Shank3:* 0.21 ± 0.005, one-way ANOVA, p = 0.05; excluding wildtype animals and comparing across mouse models of ASD, p = 0.67). This conclusion was also corroborated via pupillometry demonstrating a surprise signal during the presentation of statistically unlikely events (e.g., high contrast on the right during a leftward block) in wildtype but not mouse models of ASD (see **Fig. S2** for details).

Together, these results mimic findings from the human literature in demonstrating an attenuated use of priors^23, 24^ and the lack of a surprise signal (at least insofar as indexed via pupillometry; see^32–34^ for a similar approach) during the presentation of statistically unlikely events^21^ in mouse models of ASD. Importantly, this was a generalized finding, being true for all mouse models of ASD tested.

### Blunted accumulation of recent sensory history in mouse models of ASD

Next, we aimed at understanding the strategies employed by different animals to update their priors, and to estimate the “subjective” (vs. “objective” or experimenter-imposed) prior utilized by animals on each trial. To do so, we derived a set of models (principled and heuristic) and contrasted their performance in explaining animal behavior.

We fit a total of 10 models, comprising of variations (e.g., assumptions over symmetry of the prior, see *Methods*) over 4 broad classes. First, we simply fit psychometric curves (Fig. 2A, top row) for each of the three blocks (50:50, 80:20, and 20:80). This effort does not postulate any particular strategy animals may have employed in solving the task, but serves as a standard and adequate benchmark in accounting for responses. Second, we build a Bayesian decision maker (see *Methods*), but directly provided this model with the true prior probability of observing gratings on the left vs. right (omniscient model; Fig. 2A, second row), or allowed for a single fixed prior (fixed model; Fig. 2A, third row). These variants establish model performance when not allowing the prior to deviate from the experimentally-imposed prior, or for it to vary dynamically. Third, we use the same decision-maker as above, but also employ multiple variants of a Bayesian online change point detection algorithm (c.p. models^29, 30^, Fig. 2A, 4^th^ to 8^th^ row) to estimate the prior. The algorithm is “online” in that it only uses observations until the current trial (as opposed to the whole sequence) and is built iteratively, allowing the model to keep track of the probability that the next stimuli will be presented on the left (vs. right) with simple operations. Importantly, these models explicitly hypothesize that animals have an understanding of the task structure. For example, that there are blocks of trials wherein stimuli presentation is biased. The exact parameters estimated may of course deviate from those imposed experimentally. Lastly, we build a heuristic model (exponential weighting models, “exp. models”, Fig. 2A, 9^th^ and 10^th^) wherein animals do not know about block structures but track statistical regularities by computing a weighted average (favoring the most recent stimuli) of their recent sensory past. In a second variant of this exponential weighting model (Fig. 2A, 10^th^ row, “exp. bias model”, in red), animals in addition have a prior over counts (i.e., “pseudo-counts”) that may bias their weighted average and changes the relative weighting between observed sensory history and observation-independent a-priori counts.

**Figure 2.**
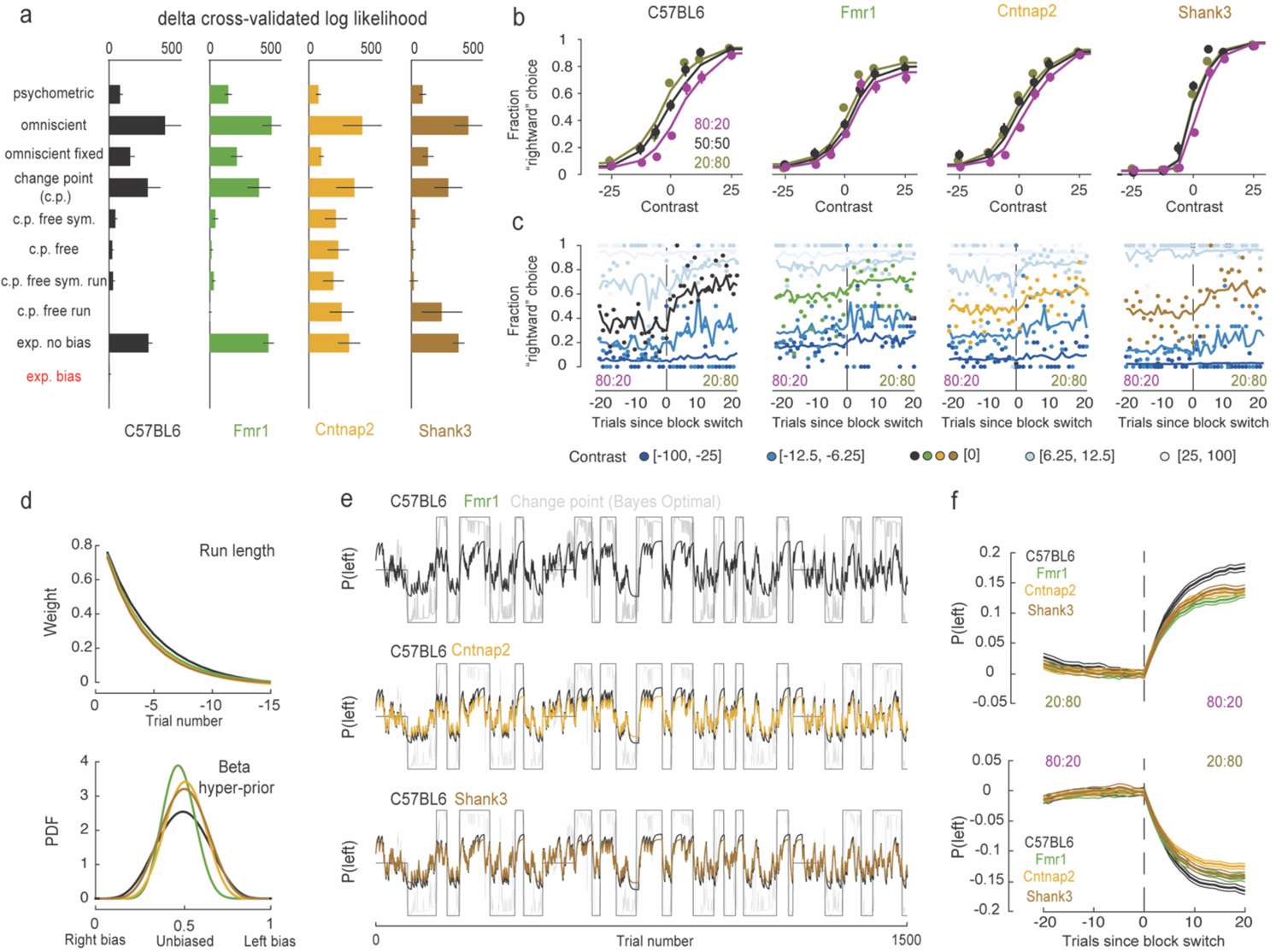
Blunted accumulation of recent sensory history in mouse models of ASD. **A.** Difference in cross-validated log likelihoods (i.e., “badness” of fit) relative to the best model (exp. bias, highlighted in red). Lower is better. Description of each model in main text. See **Figure S3** for parameter estimates from the c.p. free run model, demonstrating that while it was a comparable fit to the exp. bias model for *C57BL6* and *Fmr1* animals, its resulting parameters suggest animals did not infer the presence of experimental blocks. Color scheme for genotypes follows that of Figure 1. Error bars are ± 1 s.e.m. across animals. **B.** Fraction of rightward choices as a function of block (80:20 leftward in purple, 50:50 unbiased in black; 20:80 rightward in gold) for 4 example animals; 1 per genotype. Circles are data and lines are fits from the exp. bias model. **C.** Example fits of the exp. bias model (lines) when plotting data as a function of trials to and since block change (in this case, from 80:20 to 20:80). As expected, change in behavior is most notable for 0 contrast trials (colored). The rest of contrasts are grouped, the color gradient (from dark to light blue) following the spectrum from strong evidence for left targets to right targets. **D.** Visualization of the average exponential decay (top) and beta hyper-priors (bottom) dictating the exp. bias model for wildtype (black) and mouse models of ASD (colored). The hyper-priors being taller and narrower in ASD result in a diminished change in the subjective prior with changing environmental statistics. **E.** Illustration of an experimental sequence of trials; the experimentally-imposed probability that the stimuli will be on the left (black step functions), what an optimal observer would be able to infer (gray), and the best estimates of subjective priors for the average control animal (top; black jagged line), as well as the average Fmr1 (top; green), Cntnap2 (middle; yellow), and Shank3 (bottom; brown) animal. **F.** Subjective prior before and after block changes in the wildtype (black) and mouse models of ASD (colored). Estimates are baseline-corrected averages across animals and all transitions. Error bars represent ± 1 s.e.m.

Model comparison (Fig. 2A, based on cross-validated log-likelihood) favored the biased exponential weighted average model (“exp. bias” model, ANOVA, p = 1.46 x 10^-13^), and this was true across all genotypes (interaction term, p = 0.88). This demonstrates that we can estimate an animals’ task strategies (i.e., psychometric fits did not account best for responses, see Fig. 2B for fits from the preferred model). It also suggests that the animals did keep track of the statistical regularity embedded in the sequence of grating presentations (i.e., fixed model was not favored, see Fig. 2C for fits of the preferred model as a function of trial to and from block change), but did so via heuristics as opposed to developing a full generative understanding of the task. Lastly, it suggests that the different genotypes did not employ categorically different strategies, which allows for contrasting recovered model parameters, as well as estimate “subjective” priors.

From the biased exponential weighted average model we may estimate a time (in fact, “trial”) constant over which animals accumulate evidence in estimating the probability that the following stimuli will be presented on the left (vs. right). We may also estimate a “pseudo-count” prior, putatively biasing the count and rendering its update less dependent on observations. These are illustrated in Figure 2D and show that while the trial constant was not different across the wildtype and mouse models of ASD (top; one-way ANOVA, p = 0.15; half-width ∼ 5 trials), the hyper-priors (or “pseudo-counts”) exerted a greater influence (i.e., Beta distribution is sharper) in all mouse models of ASD relative to the control (bottom; p = 0.04). This accounts for the reduced change in choices across blocks in the mouse models of ASD (Fig. 1 and Fig. 2B **and C**). Further, when applying these parameters to a sequence of trials we can see that the subjective priors (even for the control animal) are far from the extremes imposed experimentally (0.2 to 0.8, Fig. 2E), and this effect is exacerbated in the mouse models of ASD (Fig. 2F, p < 0.003).

### A large-scale neurophysiological survey across various mouse models of ASD

Having established that multiple mouse models of ASD exhibited a computational anomaly akin to that shown by humans^21, 23, 24^ on the autism spectrum (Fig. 1) and intimated a putative computational strategy (Fig. 2), we were next interested in establishing its neural underpinning. Namely, the fact that multiple genotypes displayed the same deficit affords us the opportunity to attempt distilling causal contributions to prior coding and updating, as well as its putative dysfunction in models of ASD; i.e., what neural features are common across all mouse models of ASD and different from the control?

Answering this question requires a neural survey on the scale of the whole brain^35–37^. In turn, we used neuropixel probes^31^ (323 insertions) to record from a total of 53,219 units across 150 brain regions (Fig. 3A, see *Methods* and REF^38^ for further detail, **Fig. S4** for histology). To reliably compare neural features across genotypes we need a population of units within a defined anatomical region for all animal types. Thus, we set a threshold of a minimum of 40 units per area and genotype (2x the criteria from REF^36^). This reduced the dataset to 39,393 units across 36 brain regions (Fig. 3B, abbreviations follow the Allen Common Coordinate Framework; CCF^39^).

**Figure 3.**
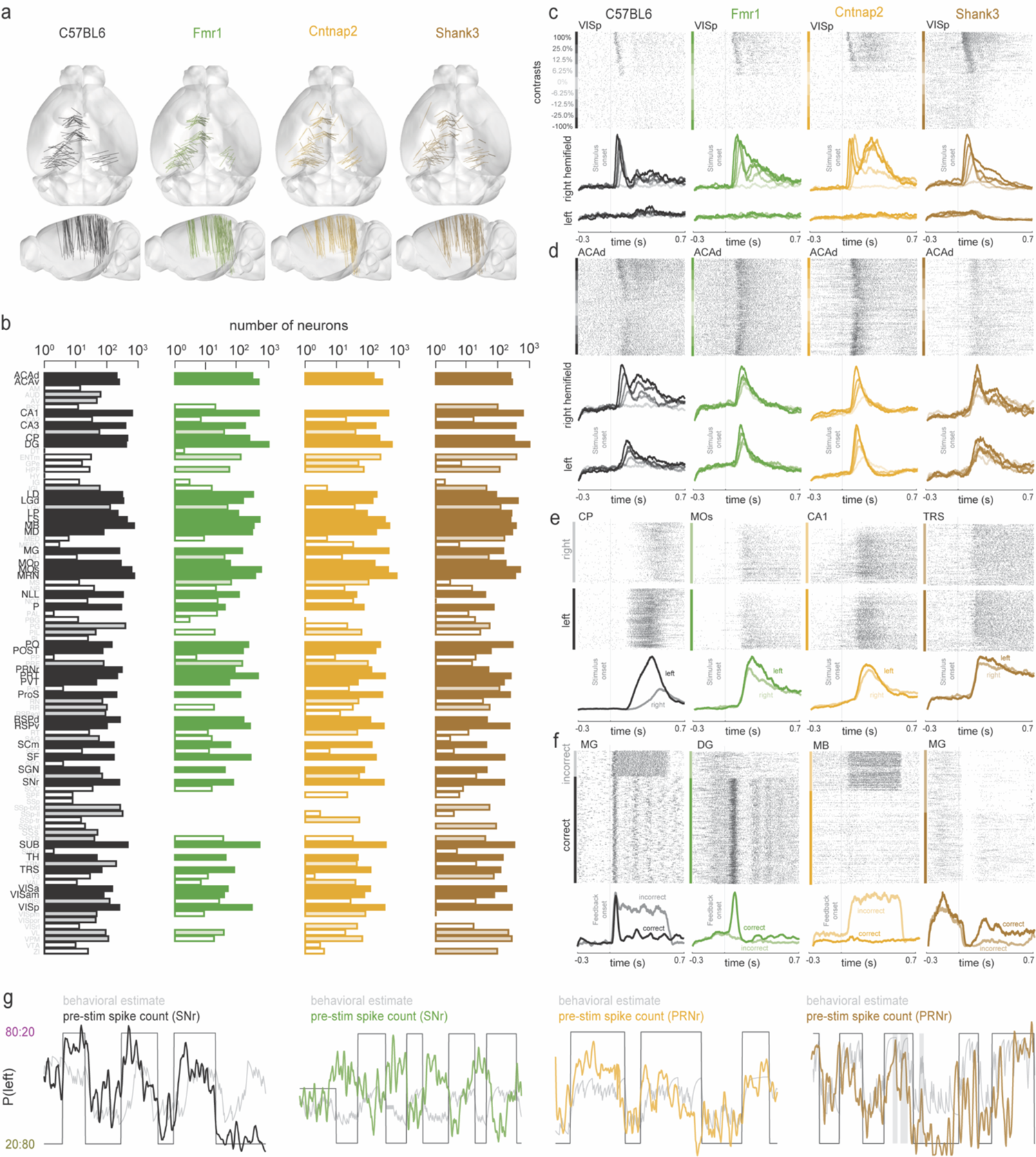
A large-scale neurophysiological survey across mouse models of ASD. **A.** Reconstruction of probe locations for the different genotypes (C57BL6 in black; Fmr1 in green; Cntnap2 in yellow; Shank3 in brown). **B.** Number of units recorded per area (y-axis; subset shown) and genotype. Bars are filled and opaque if all genotyped had at least 40 units in that area, filled and transparent if the given genotype had 40+ units but others did not, and empty if less than 40. **C.** Raster plot (top) and PSTH (bottom) to stimulus onset for example units in VISp. Raster is sorted by contrast (positive values indicating gratings presented on the right; recordings on the left hemisphere; panel **A**). **D.** Similar to **C** but showing responses in ACAd. **E.** Raster and PSTH to stimulus onset, but sorted as a function of choice. Areas are marked on the top left of rasters. **F.** Similar to **C-E,** but sorted by aligned to feedback onset and sorted as a function of correct and incorrect responses. **G.** Example spike counts (normalized; colored by genotype) before stimulus onset (−300 to −50ms) as a function of trial number (x-axis). Also plotted are the experimentally-imposed prior, and the subjective estimate of the prior for the given animal/session. In gray (rightmost panel) we highlight example periods where firing rate changes during changes in the subjective prior and stable experimental prior.

As expected, rasters and peri-stimulus time histograms demonstrated standard features of neural responses in all genotypes. Namely, visual evoked responses occurred at earlier latencies and were more robust with increasing contrast (Fig. 3C). These responses were distributed, often contra-lateral in the primary visual areas (VISp, Fig. 3C, examples shown), and regularly bi-lateral and at larger latencies elsewhere (e.g., ACAd, Fig. 3D, see Steinmetz et al., 2019). Units across many regions (Steinmetz et al., 2019; IBL et al., 2023) appeared to correlate with the choice of the animal (Fig. 3E, examples in CP, MOs, CA1, and TRS shown). Similarly, responses to feedback (e.g., incorrect response) were also distributed and could be expressed as an increase (Fig. 3F, first and third column) or decrease (third and fourth column) of firing rates (Fig. 3F. The C57BL6 and Fmr1 examples also show responses seemingly driven by the lick response). For a full characterization of these phenomena readers are referred to previous reports using very similar^35^ or identical^36^ protocols. Instead, here we focus on coding of the subjective prior, as estimated by a biased exponential weighting of recent sensory history, and on differentiating factors between wildtype mice and mouse models of ASD.

We computed spike counts within a window preceding trial onset (−300 to −50ms) and plotted these on a trial-by-trial fashion, jointly with the experimentally-imposed prior and our behavioral estimate (Fig. 2, exp. bias model) of this prior. This suggested the presence of a subset of units (Fig. 3G, 4 shown, 1 per genotype) whose pre-stimulus firing rates co-varied with block. Interestingly, close inspection showed periods where pre-stimulus firing rates changed as the subjective estimate of the prior changed, even when the experimentally imposed prior remained constant (e.g., Fig. 3G, shank3 example in brown, periods highlighted in gray). This motivates a quantitative detailing of neural responses to the subjective prior. We first take a big picture approach by examining population-level encoding (Fig. 4), and then examine unit responses and neural tuning (Figs. 5 & 6).

**Figure 4.**
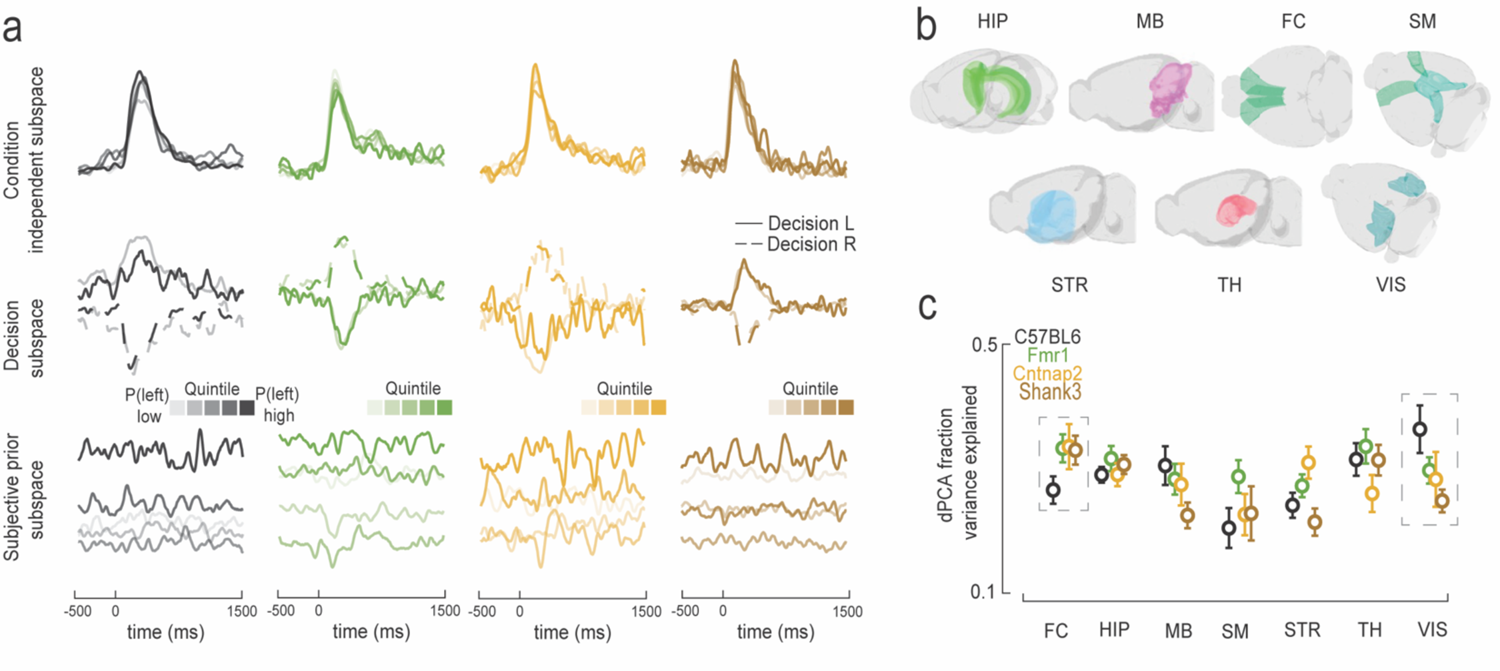
Population encoding of subjective prior shifts from visual and frontal cortices in mouse models of ASD. **A.** Four example demixed PCA (dPCA) sessions, one per genotype. Analysis was conducted to separate quintiles of the subjective prior and left vs. right decision. Top row shows a “condition-independent” subspace capturing the stimulus evoked response (only left decision – solid lines – shown for clarity). Middle row show the decision subspace appropriately separating left (solid line) and right (dashed line) choice. Bottom row shows the subjective prior subspace. Of note, the quintiles of the subjective prior are differentiated before the stimulus is presented (x-axis = 0). **B.** Categorization of brain regions into ‘macro-areas’ for statistical power and coarse summary (HIP = hippocampal areas; MB = midbrain; FC = frontal cortex; SM = somatosensory and motor; STR = striatum; TH = thalamus; VIS = visual areas; see **Table S1** for further detail on the categorization). **C.** Variance explained by the subjective prior subspace as a function of “macro-area” and mouse genotype (C57BL6 in black; Fmr1 in green; Cntnap2 in yellow; Shank3 in brown). Error bars represent ± 1 s.e.m.

**Figure 5.**
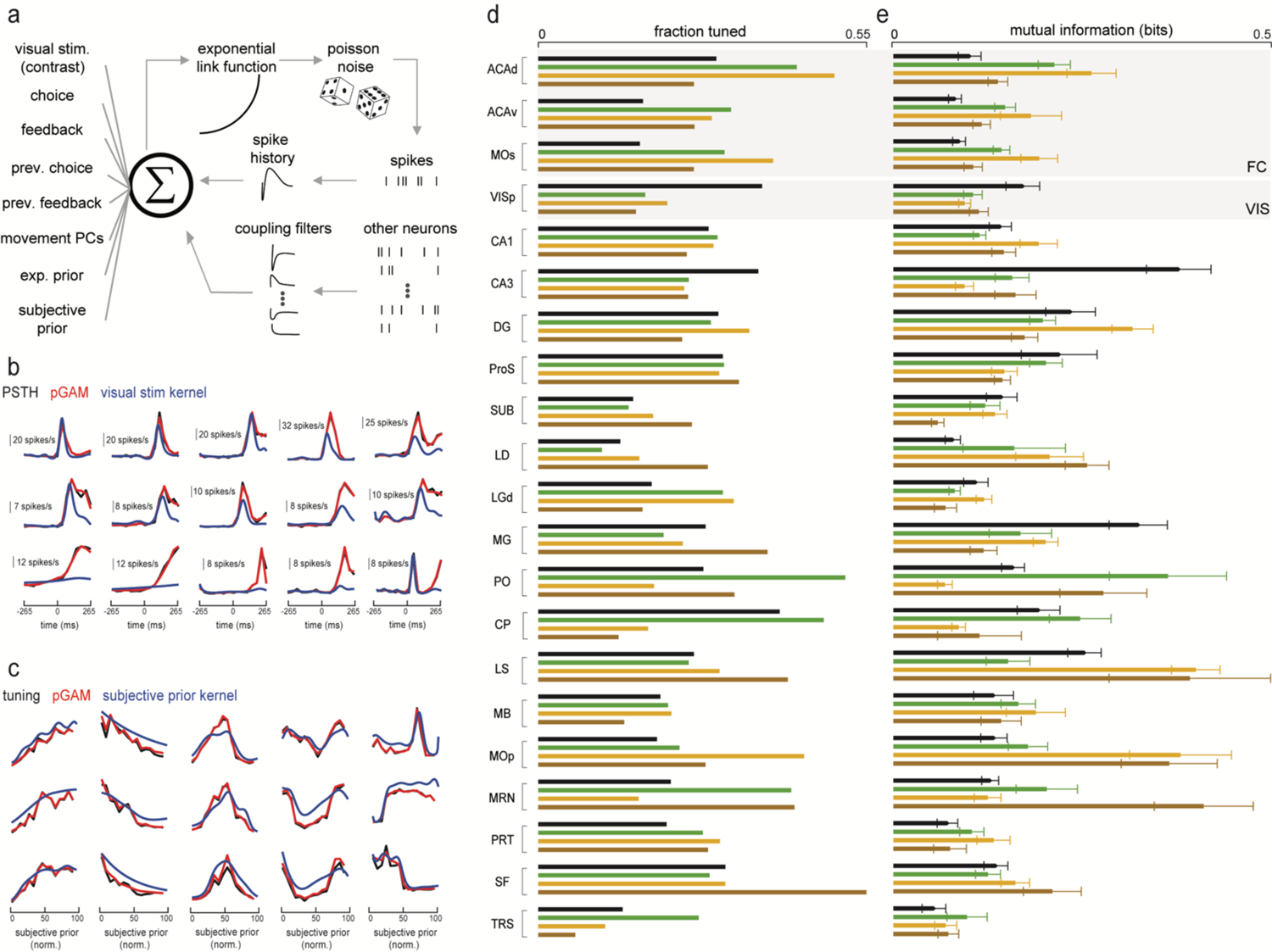
Unit encoding of subjective prior shifts from visual and frontal cortices in mouse models of ASD. **A.** Schematic of the pGAM encoding model **B.** Example units showing the empirical PSTH (black, x-axis is time), the reconstructed average from the pGAM (red), and the (exponentiated) visual stimulus kernel (blue). The latter is the contribution to the observed response the encoding model ascribed (factorized) to visual stimulus presentation. **C.** Example tuning function to the subjective prior. Follows the format from **B**, with the difference that the x-axis is not time anymore, but the value taken by the subjective prior. The x-axis is normalized such that the lowest value taken during a recording (y-axis in Figure 2B) takes a value of 0, and the maximum takes a value of 1. By definition, therefore, 0.5 corresponds to the average subjective prior of the animal. **D.** Fraction of units significantly tuned (p<0.001) to the subjective prior as a function of brain region (vertical) and genotype (C57BL6 in black; Fmr1 in green; Cntnap2 in yellow; Shank3 in brown). **E.** Follows the convention from **D,** showing the informativeness of tuning functions (measured in mutual information). Error bars are ± 1 s.e.m.

**Figure 6.**
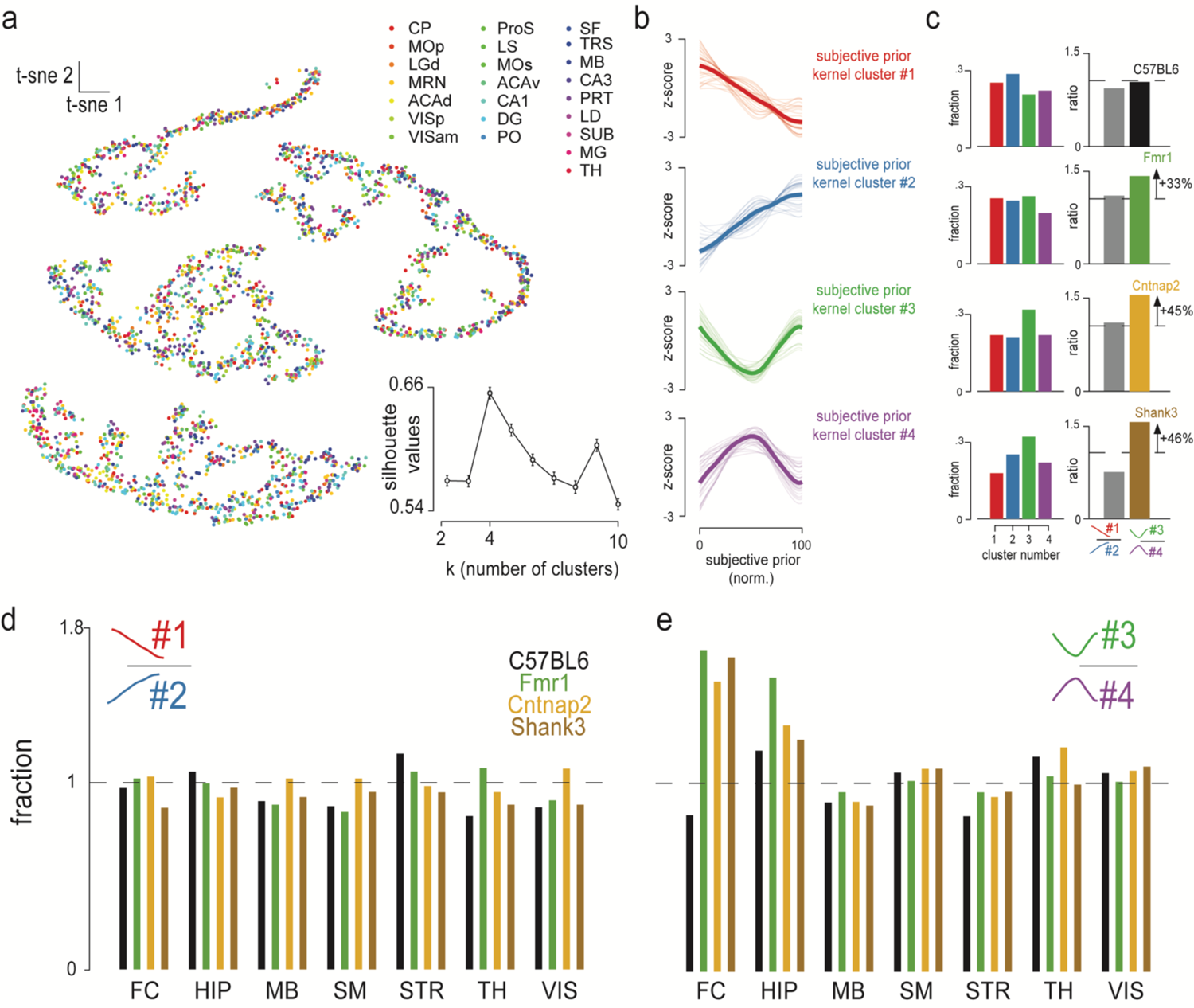
Outsized coding of deviations from long-run prior across mouse models of ASD. **A.** Two-dimensional t-Distributed Stochastic Neighbor Embedding (t-SNE) of tuning functions to the subjective prior as a function of brain area (colors). Inset shows the average and s.e.m. silhouette values (i.e., relative distance of points to others within and across clusters) as a function of number of clusters. A high silhouette value indicates that points within a cluster are well matched to their own cluster, and poorly matched to other clusters. **B.** Examples (thin and transparent) and average (dark and opaque) tuning functions to the subjective prior as a function of cluster (1 through 4). **C.** Left: Fraction of the units tuned to the subjective prior that belong to each of the 4 clusters (shown in **B**) as a function of genotype (rows; C57BL6, Fmr1, Cntnap2, and Shank3, respectively). Right: Fraction of units increasing their firing rate with decreasing (left: red) vs. increasing (left: blue) value of the subjective prior (gray), and fraction of units increasing their firing rate with increasing (left: green) vs. decreasing (left: purple) distance from the long-run prior (i.e., normalized subjective prior = 0.5). **D.** Fraction of cluster 1 / cluster 2 (gray in **C**) as a function of genotype (colors) and macro-brain area. Acronyms follow convention from Fig. 4**. E.** As **D.,** showing the fraction of cluster 3 / cluster 4. The dashed line shows a fraction = 1.

### Population encoding of subjective prior shifts from visual and frontal cortices in mouse models of ASD

We examined low-dimensional population-level encoding via demixed principal component analysis (dPCA^40^). This method is conceptually similar to PCA, but attempts to keep the learned components interpretable vis-à-vis task features (e.g., decision or prior; Fig. 4A). In the current dataset, dPCA explained 86.1% of the variance explainable by PCA (over the first 10 PCs). This analysis was performed on individually CCF-defined regions (Fig. 3B; see Fig. 4A for example sessions), yet to derive a coarse-level picture and for statistical power, we subsequently coalesced regions into ‘macro-areas’ (Fig. 4B, see **Table S1** for detail and REF^41^ for a similar approach).

This analysis demonstrated that information regarding the subjective prior (as derived from the exp. bias model above) was present throughout the brain and spanning all levels of the neural hierarchy (Fig. 4C, see^37^ for a similar finding). The total variance explained by subjective prior subspaces was 28.3% ± 0.4%, and this value was not different across genotypes (one-way ANOVA, p = 0.29). The subjective prior accounted for about two-thirds (68.0%) the variance explained by evoked visual responses (41.3% ± 0.5%), and for three times as much than decision subspaces (8.65% ± 0.1%). When splitting by brain area, we did observe differences across genotypes (ANOVA interaction term, p = 0.039), which was driven by (1) an increased population-level encoding of the subjective prior in frontal areas (ACAd, ACAv, MOs) of mouse models of ASD (variance explained; Fmr1: 31.8% ± 1.95%; Cntnap2: 32.0% ± 3.13%; Shank3: 31.5% ± 2.14%) relative to the wildtype (26.1% ± 1.94%, all p < 0.05; Fig. 4C), and (2) more prominent encoding of the prior in visual areas of the wildtype animal (34.3% ± 3.40%) relative to mouse models of ASD (Fmr1: 28.7% ± 1.79%; Cntnap2: 27.5% ± 3.73%; Shank3: 24.6% ± 1.75%, significant for wildtype vs. Shank3, p = 0.02, and showing trends for Cntnap2 and Fmr1, p = 0.10 and p = 0.11 respectively).

These results suggest that while a multitude of neural regions may show responses reflecting the subjective prior of animals^37^ the differentiating factor between animals updating their priors stereotypically vs. only modestly (mouse models of ASD) is a putative graded shift in the prior encoding from visual to frontal cortices. This analysis, however, is only coarse-level and could be driven by co-variates (e.g., correlation between prior and choice). Thus, we next examined unit properties with an encoding model disentangling the contributions of co-variates.

### Unit encoding of subjective prior shifts from visual and frontal cortices in mouse models of ASD

Spike trains were fit to a Poisson generalized additive model (pGAM^42^) including as predictors the timing and contrast of gratings, choice (left or right), and feedback (correct or incorrect). We also include the choice and feedback on the previous trial, as well as the first 10 PCs of video body-part tracking data, and the experimental (20:80, 50:50, 80:20) and subjective (exp. bias model) prior (Fig. 5A, left column). The inclusion of both the experimental prior and the immediately precedent choice/feedback ensures that units labeled as encoding for the subjective prior do not simply reflect stimulus statistics or the immediately previous choices/feedback, but in fact reflect a (weighted and biased) accumulated sensory history (i.e., the exp. bias model, see *Behavioral modeling* section). Lastly, the model attempts to also account for elements of internal neural dynamics, by allowing unit-to-unit couplings and spike-history (Fig. 5A, right column; see^43–45^ for a similar approach and **Fig. S5** for characterization of the stability of coupling filters as a function of block and genotype). Importantly, beyond capturing arbitrary non-linearities and handling co-linear predictors^46^, the specific pGAM we fit^42^ infers marginal confidence bounds for the contribution of each feature and thus allows identifying the minimal subset of factors that significantly (p<0.001) impact neural responses without computationally costly (and often unstable) model selection procedures. We achieve a goodness-of-fit (pseudo-R^2^ = 0.0745) on par with state-of-the-art machine learning techniques^47^ while reducing our encoding models by an order of magnitude (see **Fig. S6** for further model performance quantification).

We first briefly present the encoding of visual stimuli – as sanity check – before assessing the encoding of the subjective prior. In this regard, we find units that encode grating presentations throughout the brain^35, 36^ with the primary visual cortex (VISp) showing the strongest difference (in mutual information) between contra-lateral responses (arguably driven by the stimulus itself) and ipsi-lateral responses (arguably further driven by recurrent neural dynamics; **Fig. S7**). Figure 5B shows the peri-stimulus time histogram (PSTH) for a few example units, overlayed with the pGAM reconstruction and the estimated contribution of the visual stimulus kernel. Interestingly, we can observe that while for a few units (e.g., first two examples in Fig. 5B) the entirety of their response to grating presentation is (bottom-up) visually driven, for the majority of units (e.g., rest of first row and second row, Fig. 5B) their early response is sensory driven whereas their later latency responses are less so (putatively accounted for by the coupling filters). Some units showing increased firing after grating presentation were, in fact, according to the pGAM, not driven by visual stimulus at all (Fig. 5B, bottom row). Fittingly, these latter units showed a delayed evoked response relative to the “bottom-up” driven units. Overall, these results broadly align with classic characterizations of the visual pathways^48, 49^ and recent brain-wide surveys^35–36^, while also demonstrating the utility of including in the encoding model both externally-driven predictors and internal elements of neural dynamics.

The subjective prior was encoded in units throughout the brain (examples shown in Fig. 5C), in fractions (averaged across genotypes) ranging from 0.14 (Triangular nucleus of septum) to 0.36 (Anterior cingulate area; alpha set at 0.001 and thus well below these fractions; Fig. 5D). Separating by brain area and genotype, we observed that akin to the population-level results, the subjective prior was more frequently coded in frontal areas in mouse models of ASD relative to the control (coalescing ACAd, ACAv, and MOs; χ^2^ test = 30.67, p = 10^-5^; when taking the areas independently, p < 3.5×10^-4^ for ACAv and MOs, and p = 0.10 for ACAd; all mouse models of ASD > wildtype in each frontal area, with the exception of Shank3 in ACAd). Concurring with the population-level results, the subjective prior was less frequently coded in the visual cortex in mouse models of ASD relative to the control (p = 10^-5^; independently, all mouse models of ASD < wildtype, all p < 2.0×10^-4^). We also estimated the informativeness (see *Methods*) of each of these tuning functions (Fig. 5E, a continuous variable as opposed to the binary tuned vs. not tuned). These results showed greater mutual information in mouse models of ASD than the wildtype in each of the frontal areas (all p < 3.8 x 10^-3^), and reduced mutual information relative to the control in visual cortex (p = 3.04 x 10^-4^; Fig. 5E). Other areas (most prominently CA3) showed differences in frequency (p = 0.004) and informativeness (p = 8.14 x 10^-16^) of tuning in mouse model of ASD relative to the control, but these differences were either (1) not consistent within neighboring regions/established circuits (e.g., CA1 vs. CA3 or DG; Fig. 5D and **E**), (2) not true across measures (e.g., MG FT vs. MI; respectively, Fig. 5D and **E**), or (3) not observed across all mouse models of ASD tested (e.g., LS or MOp).

### Outsized coding of deviations from long-run prior across mouse models of ASD

Next, we sought to move beyond the summary characterization of frequency (Fig. 5D) or informativeness (Fig. 5E) of neural responses vis-à-vis the subjective prior, and instead examine the underlying shape of these tuning functions. We retained the top 10k tuning functions by mutual information with the subjective prior, and performed K-means clustering^50^ while allowing the number of clusters to vary from 2 to 10. We also projected the subjective prior kernels (blue in Fig. 5C) onto a two-dimensional t-sne (Fig. 6A, t-Distributed Stochastic Neighbor Embedding^51^) for visualization. The tuning functions were best described as pertaining to 4 clusters (Fig. 6A, inset shows silhouette values quantifying separateness of clusters). Cluster 1 (Fig. 6B, 1^st^ row) were units whose responses monotonically decreased as the subjective prior increased from its lowest value (normalized value of 0, p(Left) low) to its highest value (p(Left) high). In other words, these are units that fired when the subjective prior indicated a high likelihood that the next stimuli will be presented on the right. Cluster 2 (Fig. 6B, 2^nd^ row) were the mirror image of Cluster 1, with neural responses monotonically increasing from p(Left) low (norm. subjective prior ∼ 0) to p(Left) high (norm. subjective prior ∼ 1). Both these clusters code for the current value of the subjective prior. In contrast, Cluster 3 (Fig. 6B, 3^rd^ row) was comprised of units whose firing rate was minimal at a normalized subjective prior of 0.5, and fired more at both high and low values of the subjective prior. That is, these units were driven by deviations from the prior mean, the long-run prior of each animal (norm. subjective prior = 0.5). Lastly, Cluster 4 (Fig. 6B, 4^th^ row) was the inverse of Cluster 3, with units firing most readily when the subjective prior took on intermediate values. Clusters 3 and 4 code for the absolute value of the difference between the session-average subjective prior (prior mean) and the current trial subjective prior.

Units of all cluster types were present in each of the brain areas we recorded from (Fig. 6A). Thus, we examined the fraction of each cluster type throughout the brain of wildtype and mouse models of ASD. Interestingly, these were not equally distributed across all genotypes (χ^2^ test = 50.38, p < 10^-5^). Instead, Cluster 3 was over-represented in the Cntnap2 (p = 0.002) and Shank3 (p = 0.008) animals, and Cluster 4 was under-represented in Fmr1 (p = 0. 04, Fig. 6C). Clusters were uniformly represented in the wildtype (p = 0. 21). Given that Clusters 1 and 2 (coding for subjective prior value), and Clusters 3 and 4 (coding for absolute difference from the long-run prior) were mirror images of each other, we computed the relative fraction of these cluster types. This analysis showed an approximate equal fraction of units increasing or decreasing their firing rate with the value of the subjective prior (Fig. 6C, right, gray). Instead, Fmr1, Cntnap2, and Shank3 animals had, respectively, a 33%, 45%, and 46% increase in units increasing their firing rates as the subjective prior took on values further from their long-run prior (Fig. 6C, right, colored). This was not true of the wildtype animals, with an equal number of units increasing (Cluster 3) and decreasing (Cluster 4) their firing rate with deviations from their long-run prior.

When splitting across “macro-areas” (**Table S1**), it remained true that across genotypes and areas a similar fraction of units either increasing or decreasing their firing rate with the value of the subjective prior (Fig. 6D, χ^2^ test, all p > 0.13). That is, across areas, wildtype and mouse models of ASD coded similarly for the instantaneous value of their prior. Instead, the outsized population of units coding for deviations from the prior mean in ASD (Fig. 6C) seemed to be driven by frontal cortex (χ^2^ test = 113.14, p < 10^-5^), and to a lesser extent by the hippocampal formation (χ^2^ test = 17.40, p = 5.85 x 10^-4^; Fig. 6E).

Together, the results demonstrate both global and local differences in unit coding of the prior across animals flexibly and inflexibly updating their expectations. Globally, results suggest that relative to wildtype animals, mouse models of ASD more heavily rely on frontal cortices, and less on visual cortices, to encode their priors. Locally, the tuning functions in frontal cortex demonstrate a selective preponderance of units increasing their firing rates with deviations from the long-run prior of the animals.

### Lack of sensory-driven prediction errors in frontal cortex of mouse models of ASD

The updating of priors ought to be driven by the observation of statistically unlikely events (i.e., the accumulation of these events ultimately results in an update of what is considered statistically likely). Thus, we examined how the encoding of gratings was modulated by their statistical likelihood. To do so, we fit the pGAMs^42^ separately for each experimental block (80:20 and 20:80) and compute mutual information (see *Methods*) at each contrast. We observe that when most stimuli were presented on the left/right hemifield, the encoding of stimuli (as index by mutual information) presented on the opposite side was strengthened (Fig. 7, all brain areas and genotypes combined, p = 1.27 x 10^-43^; 20:80 > 80:20 for negative contrasts, and 20:80 < 80:20 for positive contrasts). This is in line with the theoretical framework of predictive coding^52^.

**Figure 7.**
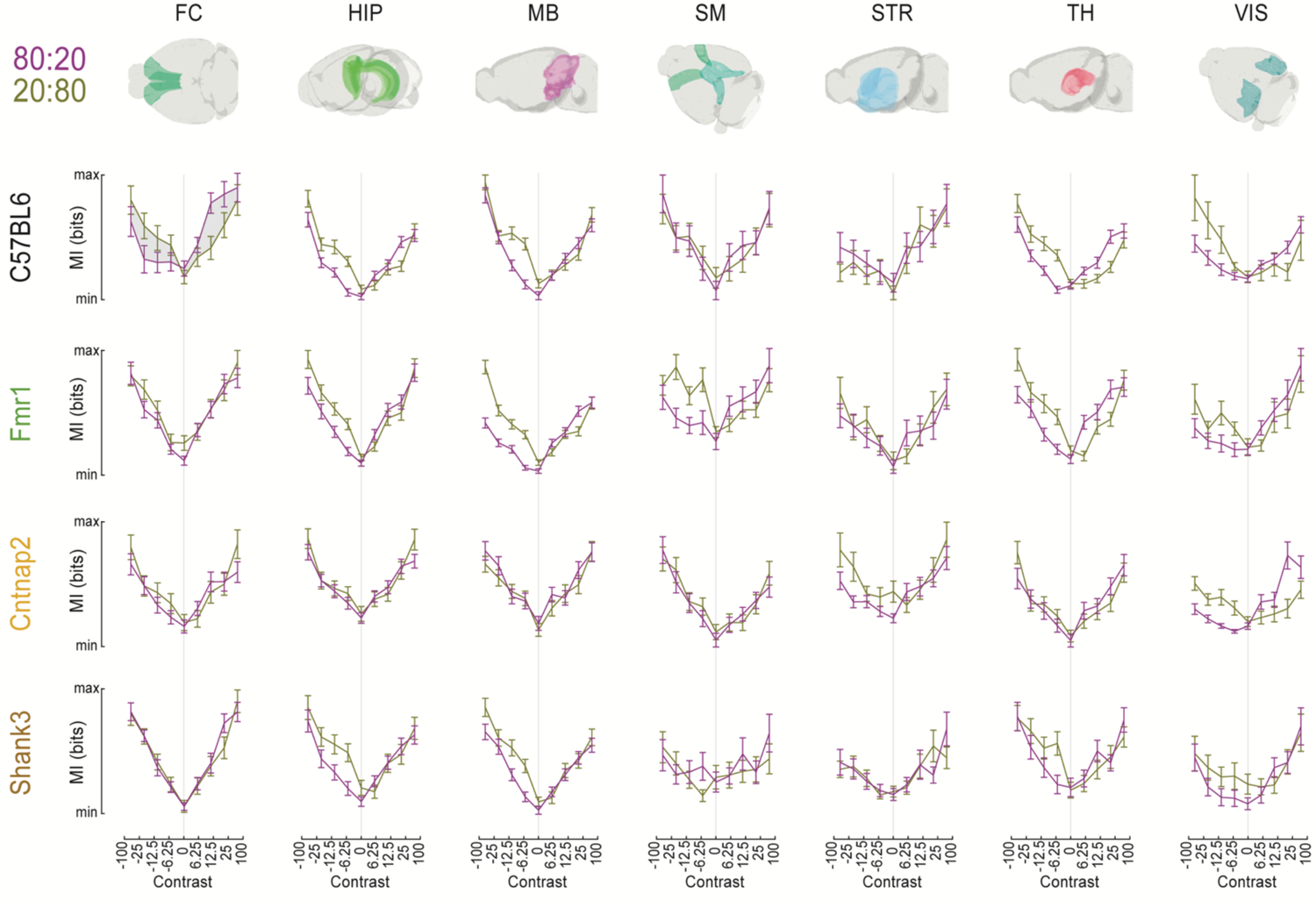
Lack of sensory-driven prediction errors in frontal cortex of mouse models of ASD. Mutual information (MI, y-axis) as a function grating contrast (y-axis; negative contrasts are gratings presented on the left hemifield), experimental block (leftward bias block in purple, 80:20), macro-area (columns), and genotype (rows). Error bars are ± 1 s.e.m.

We observed abundant variability when separating across “macro-areas” (**Table S1**) and genotypes. For instance, the Cntnap2 animals showed a somewhat widespread lack of prediction errors (e.g., HIP, MB, SM, ANOVA interaction term, all p > 0.11) which was not evident in the other genetic models of ASD (see HIP, MB, SM in Fmr1 & Shank3, all p < 0.05). Similarly, across a number of brain regions (e.g., TH, VIS) we observed the presence of sensory-driven prediction errors in control animals, and not in a subset of the mouse models of ASD (e.g., TH in Cntnap2 and VIS in Shank3). Most strikingly, it was only in frontal cortices where we observed a differential coding of expected and unexpected stimuli in control animals (ANOVA interaction term, p < 0.001) and the lack thereof across all mouse models of ASD (all p > 0.61; Fig. 7, first column). This lack of sensory-driven prediction errors in frontal cortex of mouse models of ASD is also observable in grand-average PSTHs (**Fig. S8**).

## Discussion

We examine the behavioral, computational, and neural underpinnings of prior updating in wildtype and three different monogenic mouse models of ASD (Fmr1, Cntnap2, and Shank3). Behaviorally, the results show that all genotypes linked to ASD under-utilized the statistical regularities present in their environment (see^53^ for a similar finding in the Cntnap2 rat model). These animals were also less surprised – at least insofar as indexed by pupil diameter – during the presentation of statistically unlikely events. Computationally, behavioral modeling suggested that the mouse models of ASD and the wildtype animals used a similar task strategy. Namely, the animals did not learn a “generative model” of the task, with, for instance, the location of targets changing within blocks of a minimum length. Instead, they performed a weighted and biased average of recent sensory observations. The animals inflexibly updating their expectations had a stronger (observation-independent) hyper-prior over the predicted target locations (i.e., “pseudo-counts”). This leads to a blunted update of (observation-dependent) priors. Neurally, we show both at the population and unit levels a widespread coding of the subjective prior^37^, with a shifting in the balance of this encoding from sensory to frontal cortices in ASD. We also observed local changes with frontal cortices (ACAd, ACAv, and MOs). That is, while mouse models of ASD did not differ from the wildtype regarding their coding of the instantaneous prior, they did show an outsized presence of units coding for deviations from the animals’ long-run prior (see^54^ for a similar distinction between “immediate” and “second-order” priors and for evidence showing that the orbitofrontal cortex encodes the latter). Lastly, we demonstrate that neural responses to unexpected observations – precisely those that should lead to an update of our internal models – were augmented relative to expected stimuli in frontal cortices of wildtype animals, but not in mouse models of ASD.

Overall, the results suggest an under-weighting of sensory observations vis-à-vis a-priori expectations in ASD – both via global (i.e., over-weighting frontal vs. sensory cortices) and local (i.e., under-weighting of unexpected stimuli in frontal cortices) mechanisms. This is largely consistent with theoretical accounts suggesting an over-estimation of sensory volatility^21^ or inflexible predictions^20^ in humans on the autism spectrum. However, our results also suppose important aspects in which these theories ought to be revised.

The volatility accounts of ASD^21^ are largely rooted in a hierarchical predictive process (e.g., the hierarchical Gaussian Filter^55, 56^) with successive levels coding for beliefs about (1) the stimulus, (2) the probabilistic nature of the task (e.g., 80:20 vs. 20:80), and (3) the dynamics of this probabilistic association (i.e., volatility). However, the behavioral modeling presented here suggests that, at least in the current context, animals do not build a generative understanding of the task and thus do not have an explicit representation of volatility. It may be argued that while mice are not capable of these “deep” hierarchical inferences, humans are, and thus this is a species difference. However, modeling of human responses in a similar task^30^ suggests that humans also performed a weighted and biased average of recent sensory observations, as opposed to full inversion of a generative model. Similarly, simulations demonstrate that explicit inferences over volatility are not needed in explaining human behavior in volatile environments. In fact, models positing an explicit representation of volatility perform worse than models simply accounting for low-level uncertainty^57^. Lastly, it is interesting to note that a contrast of BOLD correlates of hierarchical predictive processes in neurotypical and autistic individuals showed strongest differences at “intermediate” levels of this hierarchy, and not in the representation of volatility. This “intermediate” level of the hierarchy reflects the probabilistic nature of the task (here, 80:20 vs. 20:80) and was linked to activity in the anterior cingulate cortex^15^ – very much in line with our findings indicating a key role for frontal cortices (including the anterior cingulate) in engendering the inflexible update of priors in ASD.

The inflexible predictions account of autism^20^ suggests a “high and inflexible” weight attributed to prediction errors in ASD. Our results demonstrate not the inflexible nature of these prediction errors, but their total absence in frontal cortices in mouse models of the disorder. Further, we observed a preponderance of units coding for deviations from the prior mean in mouse models of ASD. To the best of our knowledge, these units – not coding for the immediate prior, but for its deviance from the prior mean – are not part of current neuroscience or computational psychiatry theory, and ought to be incorporated into accounts of prior updating and its anomaly in ASD. Fitting, the anterior cingulate – where we observe an abundance of these units – is well established as an error monitoring node^58^ and is causally involved in engendering prediction errors in lower-level sensory areas (e.g., primary visual cortex^59, 60^). In future work it will be important to simultaneously record from ACAd/ACAv/MOs and visual cortices to further understand the relation between units coding for deviations from the animals’ long-run prior, and prediction errors in both frontal and sensory cortices. Similarly, in future work it will be interesting to discretize behavior into periods defined by different task strategies or internal states, and examine how accumulation not only of sensory history, but also action history^37^, may influence prior updating in each of these periods (e.g., engaged or disengaged^61^) and across genotypes. Indeed, while here we focused on sensory history as a mechanism to build expectations, it is possible that particularly when animals oscillate between engaged and disengaged states, action history better accounts for perseverative states^37^.

In conclusion, here we uncover a common computational and neural anomaly across distinct genetic mouse models of autism. The computational deficit mimics recent behavioral findings from human computational psychiatry^21, 23, 24^ and thus supposes an exciting translational opportunity to further understand the neurobiology of ASD. These results demonstrate a degree of biological degeneracy^62^ wherein different genetic perturbation may lead to similar neurophysiological consequences, both at a brain-wide scale and within local populations in frontal cortex.

## Acknowledgements

The authors express their gratitude to all IBL staff, and in particular to Steven J. West for histology and Matthew R. Whiteway for video tracking. We also thank Miranta Louka and Janna Aarse for help and expertise with animal behavior and testing. We thank Alejandro Pan-Vazquez for thoughtful reading and editing of this work. This work was supported by grants from the Wellcome Trust (216324), the Simons Foundation, and The National Institutes of Health (NIH U19NS12371601). J.P.N. was supported by NIH K99NS128075.

## Author contribution

Conceptualization: J.P.N, D.E.A; Data Curation: J.P.N., E.B., J.B..; Formal Analysis: J.P.N., E.B.; Funding Acquisition: J.P.N., D.E.A.; Investigation: J.P.N., J.B.; Methodology: J.P.N., J.B.; Project Administration: C.S., D.E.A.; Resources: L.A..; Software: J.P.N., E.B., L.A.; Supervision: D.E.A.; Validation: J.P.N., E.B., J.B.; Visualization: J.P.N., E.B., L.A.; Writing – original draft: J.P.N.; Writing – review & editing: J.P.N., E.B., L.A., C.S., D.E.A.

## Online Methods

### Animals

Experiments were performed in a total of 75 male and female mice of mixed genetic background (C57BL/6j), between ∼10 (headbar implantation and initial training) and ∼29 weeks of age (max number of sessions across all animals = 97). On average animals were 22.2 weeks old during neurophysiological recordings. In addition to wildtype control animals (n = 15), three different monogenetic mouse models of ASD were used. *Fmr1 KO*^25^ (male are *Fmr1^-/y^,* female are *Fmr1^-/-^*, no sex difference, n = 19; JAX 003025) mice have a neomycin resistance cassette replacing exon 5 of the fragile X mental retardation syndrome 1. *Cntnap2^tlacZ/tlacZ^*(n = 21; JAX 028635) mice^26^ have a dysfunctional contactin associated protein-like 2 gene by replacement of the exon 1 on the Cntnap*2* gene. *Shank3B^+/-^* (n = 20; JAX 017688) mice^27^ have a neocasette replacing the PDZ domain (exons 13-16) on the Shank3 gene, resulting in altered expression of the synaptic scaffolding protein expressed in the post-synaptic density of excitatory synapses. These animals were used as they are well-established models of ASD (REF^63, 64^; relevance to humans^65^), have previously been used in attempts to establish commonalities across mouse models of ASD^66–68^, and are visually similar to wildtype animals sharing mixed genetic background. The latter point is important given that behavioral training and testing were conducted by hypothesis-naïve experimenter (J.B.). Male and female mice of the same genotype were first analyzed separately to assess potential sex-related differences in behaviors. Given no differences were observed, male and female mice of the same genotype were grouped together for final analyses. All procedures performed in this study were approved the Institutional Animal Care and Use Committee (IACUC) at New York University.

### Surgeries

Each animal had two surgeries. A first one to secure a headbar on their skull allowing for head fixation, and a second one to perform craniotomies allowing for neurophysiological probe insertions.

For headbar implants, mice (∼10 weeks old) were initially anesthetized by placing them in an induction box at 3-5% isoflorane. They were then fixed in a stereotaxic frame and maintained anesthetized at 1-1.5% isoflorane. Under a microscope (M60, Leica), the dorsal surface of the skull was cleared of skin and periosteum, bregma and lambda were marked, the lateral and middle tendons were removed using fine forceps, and the headbar was placed and cemented on a levelled skull. A small amount of cyanoacrylate (VetBond; World Precision Instruments) was applied to the edges of the skin wound to seal it off and avoid future infections. Finally, the exposed skull was covered with clear UV-curing optical glue (Norland Optical Adhesives #81; Norland Products).

On the first day of neural recordings up to 4 micro-craniotomies were made, either with a dental drill or a biopsy punch. A gold pin touching the brain was implanted for referencing. The induction procedures followed that of the headbar implants, and craniotomies targeted −2.7mm ML/-3.5mm AP, −1.76mm ML/-2.00mm AP, −0.40mm ML/-1.06mm AP, and −0.80mm ML/0.50mm AP (negative ML values indicating the left hemisphere, negative AP values indicating posterior to Bregma). Craniotomies were covered with a low viscosity silicon sealant (Kwik-Cast, World Precision Instruments) to prevent drying. Animals were given at least 4 hours of recovery before neural recordings.

### Behavioral procedures

Following headbar implantation, animals were given at least 3 days of recovery. Then, they were handled for at least 15 minutes/day for 2 days. On the second day, the mouse was allowed to explore the behavioral rig for 10 minutes. The following three days (20, 40, and 60 minutes, respectively) the mice were head-fixed and passively presented with full-contrast (100%) Gabors (vertical orientation, 1/10^th^ of a cycle per visual degree masked by a Gaussian window of 7 degrees). The wheel used to make responses was locked. The gratings appeared on either the left or right visual field (35 degrees eccentricity and 0 degrees elevation) for an average of 10 seconds. The Gabor then moves to the center of the visual field (0 degrees eccentricity) for 1 second. The animal is given reward (3 micro-liters, 10% sucrose) 500ms after the presentation of the Gabor in the center.

Active training began on the 4^th^ day of head-fixation. The wheel was unlocked and in closed-loop with the Gabor. Gabors are presented given that the animal did not move the wheel (<2 degrees) during a quiescence period (exponential distribution, 200-500ms range, 350ms average). Initially, Gabors of either 100% or 50% contrast were presented (left or right, equal probability). A tone (100ms duration, 10ms ramp, 5kHz) was played at Gabor onset. If the mouse moves the Gabor to the center of the screen within 60 seconds of presentation, it is rewarded (3 micro-liters). If it moves the Gabor in the opposite direction by a similar displacement (35 degrees), the trial is considered incorrect. If it does not respond within 60 seconds, the trial is timed-out. In either of the latter two cases a noise burst is played for 500ms. At the onset of this active phase of training, the Gabor moves 8 visual degrees per millimeter of movement at the wheel surface. If the mouse completes at least 200 correct trials within a session (typically ∼45mins) the gain of the wheel for all future sessions is halved, remaining at 4 visual degrees/1 millimeter of movement at the surface of the wheel. Similarly, if a mouse completed 200 trials in the previous session, the reward volume was lowered by 0.1 micro-liters until a floor of 1.5 micro-liters was reached.

Behavioral training on this unbiased version of the task (i.e., probability left vs. right visual stimuli = 50:50) had six phases. First, only 100% and 50% Gabors were presented. If the animal performed above 80% correct, it moved to phase 2, where 25% contrast Gabor was added to the set. Similarly, if the animal performed above 80% correct, it moved to phase 3. In this phase, 12.5% contrast Gabors were added to the mix. To progress to phase 4, animals had to complete 200 trials within a session, regardless of performance. In phase 5 the 0% contrast was added. If animals completed 200 trials within a session, regardless of performance, they advanced to phase 6. In this last phase the 50% contrast was dropped. The mice were considered to be trained on this unbiased visual detection task if they were on phase 6 and completed at least 200 trials performing above 80% correct for 100% contrast trials for 3 consecutive sessions. Further, their psychometric estimates for the combined last 3 days had to be; bias below 16, threshold below 19, and lapses below 0.2. If animals did not learn this task within 40 sessions, they were considered untrainable. A small minority of animals were trained for over 40 sessions due to long breaks (2+ weeks) in training at an early stage.

Animals who successfully trained on the unbiased version of the task, were moved to a “biased” version of the task, wherein animals must use a dynamically updating prior in order to reach optimal performance. Namely, each session started with 90 trials where the Gabors appeared with equal probability on the left and right visual field. The side (and thus correct response) for 0% contrast Gabors was chosen randomly. After these initial 90 trials, stimuli were presented in blocks. In one block, Gabors are presented on the left with probability 80% (right, 20%). In the other block type, Gabors are presented on the left with a probability 20% (i.e., 20:80). In a given session, there was an equal probability that the first biased block were “leftward” or “rightward”. Gabors of 0% contrast are rewarded according to the prior. The number of trials for each biased block is drawn from an exponential distribution with a mean of 60, a minimum of 20 and a maximum of 100 trials. This yields an almost flat hazard rate (corrupted by the clipping of a maximum trial number). Importantly, the change in blocks is not signaled. Animals performed 10-25 sessions of this biased task before being moved to physiology (no requirement on performance, at difference from^36, 38^).

### Neural recordings

Neural recordings were performed with Neuropixels 1.0^31^ (Imec; Belgium) on the acquisition configuration (AP: 30kHz, gain = 500; LFP: 250Hz, gain = 250) recording from the bottom 384 sites of a 1-cm shank. Probes were mounted on a steel rod which was in turn held by a micro-manipulator (uMP-4; Sensapex Inc.). Probes had a soldered connection to short the external reference to ground, and this latter one was connected to a gold pin fixed on the skull and in contact with the brain. Silicone artificial dura repair compound (Dura-Gel; Cambridge NeuroTech) was placed over the craniotomies during recordings. Prior to insertion, probes were labeled for subsequent histological reconstruction (see below). On most recording days, we inserted two probes. One per craniotomy. The probes were lowered into position at approximately 10 µm s^−1^. Electrodes were allowed to settle for ∼10 minutes before starting the recording. Data were acquired via a PXIe (PXI-1000; National Instrument) using SpikeGLX (Janelia Research Campus) and stored on a PC and cloud for subsequent analyses. At most, over 4 consecutive days we performed 8 insertions in each animal; 2 per craniotomy (4 craniotomies), one medially and one laterally at 15 degrees angle from vertical. In between recordings days we covered craniotomies and exposed skull with silicon sealant (Kwik-Cast, World Precision Instruments).

### Probe labeling

For histological reconstruction, we labeled the probes with CM-Dil (Thermofisher V22888) immediately prior to insertion. Neuropixels were secured onto a micromanipulator and lowered under a microscope (M60, Leica) onto a coverslip or parafilm containing the dye (1uL). The tip of the probes were maintained in CM-Dil until the dye dried out (∼20 seconds).

### Histology and probe reconstruction

We followed the procedures standardized by the International Brain Lab^38^. Namely, mice were given a terminal dose of pentobarbital within the peritoneal cavity. Then, PBS followed by a 4% formaldehyde solution (Thermofisher 28908) in 0.1M PB pH 7.4 were perfused through the left ventricle. The brain was subsequently dissected and post-fixed in formaldehyde for a minimum of 24h at room temperature. The tissue was then washed and stored for up to ∼5 weeks in PBS at 4°C. The brains were then embedded in a 5% agarose gel block and imaged via a serial section two-photon microscopy (whole brain coronal image stacks acquired at a resolution of 4.4 x 4.4 x 25.0μm) under control of custom software (*BakingTray*). Image tiles were then assembled into 2D planes (*StitchIt*), down-sampled to 25μm isotropic voxels, and registered to the adult mouse Allen Common Coordinate Framework (CCF) using *BrainRegister*, an elastix-based registration pipeline with parameters optimized for mouse brain registration. Next, we reconstructed the location of probes by manually tracing the florescent dye on CCF-aligned coronal and sagittal images using a Python-based image viewer (*Lasagna*; Campbell et al., 2020) equipped with a plugin tailored for this task. Lastly, we manually aligned electrophysiological and anatomical landmarks along the probe trajectory using a custom tool (see^38^ and associated protocols for further detail).

### Spike sorting and curation

Data were spike sorted with Kilosort 2 (KS2^69, 70^), and/or a custom python version of the algorithm. Entire sessions were then inspected by constructing “drift maps” (channel x time, spikes as dots). If significant drift was evident, the session was rejected. Data were then manually curated via visual inspection (e.g., waveforms, auto-correlograms) with the Phy graphical user interface. Units were included in the analyzed dataset if (1) their average firing rate was over 0.5 Hz, (2) the automated labeling of units by KS2 indicated the unit as “good” (i.e., single-cell), (3) during manual curation the unit was not labeled as “noise”, and (4) it had a presence ratio (1 minus the fraction of 1 minute bins in with no spikes) above 0.9. Throughout the report we refer to the clusters satisfying these criteria as “units” given the possibility that a subset of these clusters were multi-unit, even if labeled as single units by KS2.

### Video recordings and pupil tracking during neurophysiology

We briefly describe the video analysis pipeline, which is fully detailed elsewhere^71^. We recorded videos (CM3-U3-13Y3M-CS, Point Grey) of the animals performing the task from 3 cameras/angles: top, right, and left (**Fig. S2A** shows the left camera). In the current analysis, we used the left (60hz, 1280 x 1024) and right (150hz, 640 x 512) cameras, with the latter being flipped and spatially up-sampled to resemble the left camera. We detect 4 regions of interest (ROI) on each frame (**Fig. S2A,** inset, red rectangles), crop these, and apply a separate neural network to each ROI to track features of interest. In addition to other body parts we tracked the top, bottom, left, and right corners of the pupil via *DeepLabCut*. Pupil tracking was not reliable on a number of sessions, likely due in big part to the fact that the camera positions were not optimized for pupil tracking but for simultaneous paw and tongue tracking. Thus, we performed pupillometry analyses only on the subset of sessions (n = 58 sessions) with reliable pupil diameter estimation. In each session, we dropped frames with a likelihood < 0.9 and smoothed the pupil diameter estimation. We then averaged pupil diameters across the left and right cameras, and subsequently across animals within a given genotype. We consider pupil responses to differentiate 80:20 and 20:80 blocks if p < 0.05 for at least 10 consecutive samples (see^72, 73^).

### Modeling and analyses Behavioral analyses

Training times (Fig. 1C) were determined as the number of sessions *t* (typically 5 session per week) for an animal to reach 80% correct on trials with contrast 100% or 50% (“easy trials”). To account for infrequent and sudden drops in performance (likely driven by stress caused during head-fixation), the fraction of correct trials on “easy trials” as a function of session was computed within a moving average of 3 sessions (window centered on session *t*). Psychometric curves (Fig. 1D) were fit by a parametric error function (**Eq. 1**, see^36^) with four free parameters; bias (μ), threshold (σ), and lapse rates for stimuli presented on the left and right (respectively, γ and λ). These functions fit the data well (R^2^: 0.99 ± 9.18 x 10^-4^) and equally across genotypes (p = 0.91). For visualization (Fig. 1F), we construct an average psychometric curve by prior across all animals of a given genotype, and then these are subtracted.

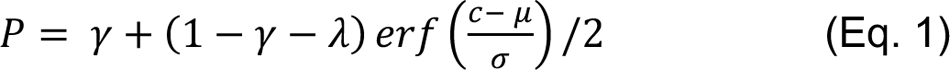

### Behavioral modeling

We build an ideal observer that has access to a noisy representation of the current stimulus (stimulus *s*, noisy measurement *x*) and the unbiased history of category labels (*C*_1:*t*_ experienced up to the present moment. We can assume an unbiased history of category labels given that animals are given feedback on each trial. The observer does not have access to the entire sequence of stimuli presented in a session, but only until trial *t*. Further, we aim to build an iterative formalism, which allows the observer to perform online inference in a tractable manner (i.e., tracking sufficient statistics and performing a simple operation at each new observation). The observer model can be decoupled into two components, a prior-tracking model computing an a-priori probability of observing a given stimulus (e.g., left) given previous observations (*C*_1:*t*_, and a perceptual decision-making model at trial *t*. We detail each of these in turn.

#### Prior Tracking

##### Online change-point detection

The Bayesian change-point detection model estimates the posterior distribution over the current run length (*r*_t_, i.e., number of trials since the last change in block) and category probabilities (i.e., left or right) given the data so far observed (category labels until trial *t*, *C*_1:*t*_ see^29, 30^). The predicted probability that the next trial presented will be on the left is,

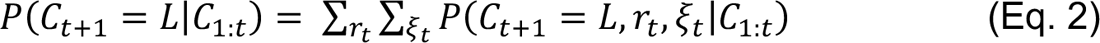

where ξ_t_ is the state of the previous block (ξ*_t_* ∈ *S*_π_, with *S*_π_ = {0.2, 0.5, 0.8}). This predictive distribution over future category may be decomposed as,

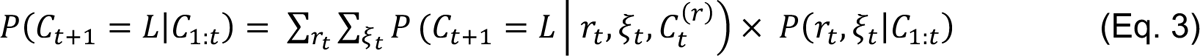

with *C*^t^ being the category labels associated with the run length *r*_t_, and the second term of **Eq. 3** being the run length and block posterior. Via iterative expansion, we can write,

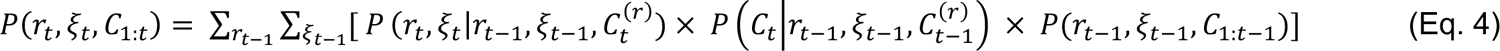

Of note, **Eq. 4** is recursive (last term being the joint distribution from the previous iteration) and affords computing *P*(*r_t_*, ξ*_t_*|*C*_1:*t*_), given that,

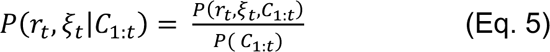

The second term in **Eq. 4** (and the first of **Eq. 3**) may be computed as,

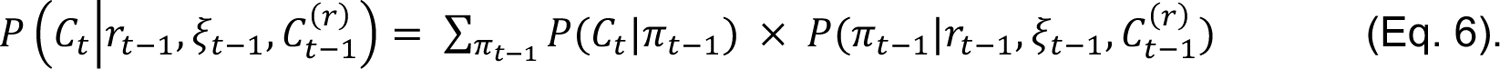

In turn,

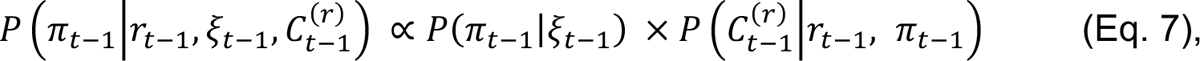

which are respectively the transition probability between blocks in the task (which is experimentally imposed) and the sequence likelihood. By definition, the transition matrix between blocks is,

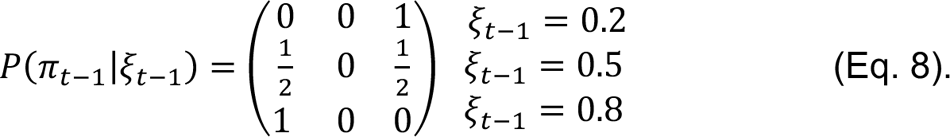

And the sequence likelihood is,

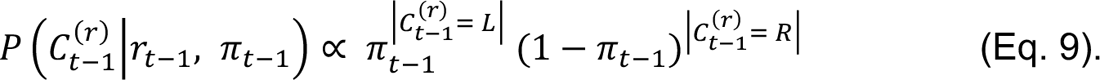

Returning to **Eq. 6**, we thus have,

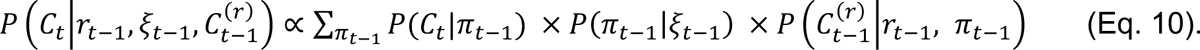

Finally, returning to **Eq. 4**, we are missing the first term. We can write,

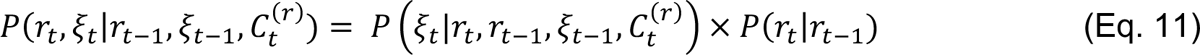

where the terms are respectively the previous-block update and the run-length update. Both of these are again experimentally imposed, and defined as,

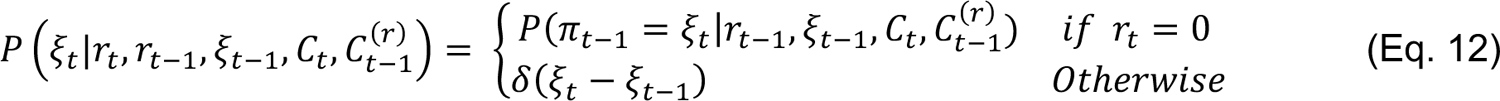

and,

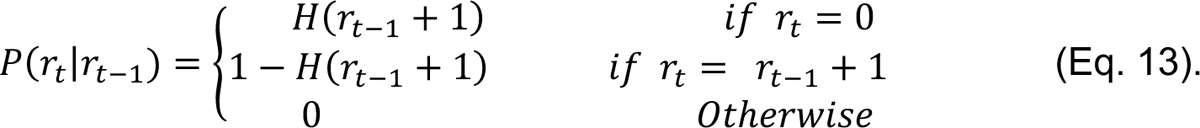

*H* is the hazard function, the instantaneous probability on each trial that there is a change-point. Namely,

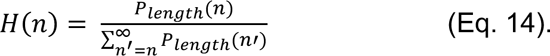

In the particular case of this experiment where block lengths are drawn from a truncated decaying exponential with time constant 60 and minimum and maximum blocks respectively of 20 and 100 trials, we have,

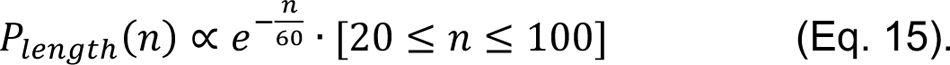

This hazard rate is approximately constant, until *n* becomes large (∼ 80 trials).

##### Exponential weighting

The exponential-averaging model postulates that animals do not explicitly know about the task structure (e.g., possessing blocks wherein stimuli presentation is biased on one side). Instead, it computes a smooth estimate of category probability by taking a weighted average of previously experienced category labels, giving more weight to recently experienced labels:

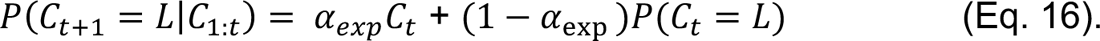

The time constant of memory decay for this model is,

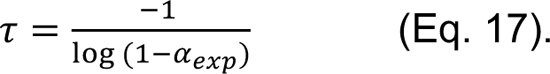

We account for conservatism^74^ in a biased version of this exponential weighting model; the exponential weighted average model with bias (exp. bias model, best model in main text). In the latter, the estimates from **Eq. 16** are biased by adding “pseudo-counts” to the observations of “left” and “right” in past trials. This is equivalent to placing a Beta distribution hyper-prior with parameters α and β on the estimated probability from **Eq. 16** (see^30^ *ExpBias* model for detail).

##### Perceptual decision-making model

The stimulus, *s*, on each trial is drawn from a set of contrasts {-1, −0.25, −0.125, −0.0625, 0, 0.0625, 0.125, 0.25, 1}, where negative values indicate stimuli presented on the left visual field. The observer does not have direct access over these stimuli, but only to a noisy measurement, *x*. We assume,

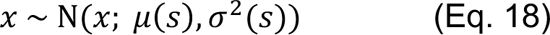

where Ν defines a Gaussian distribution, and μ and σ^2^ are respectively means and variances that depend on the stimuli presented, *s*. The ratio of posterior probabilities that a stimulus was presented on the left and right, or the decision variable, is

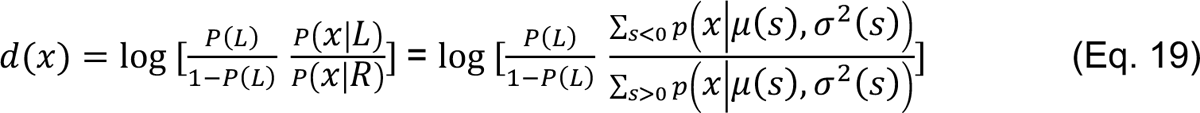

where *P*(*L*) is computed according to the prior-tracking model of choice (above) and we assume that a response “Left” is made if d(x) > 0, and “Right” otherwise. Lastly, in the case the animals do not lapse, we can write.

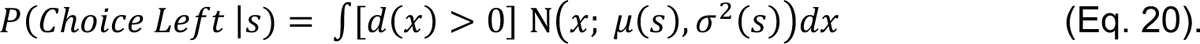

Or if the observer has an unbiased non-zero lapse rate, we can write,

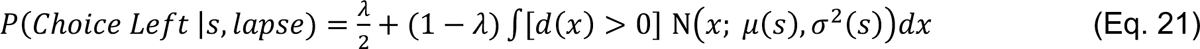

where λ is the probability that the mouse responds randomly.

##### Model fitting and comparison

We fit 10 models (Fig. 2A) to the responses of each subject. The *psychometric* model is descriptive, not attempting to explain internal representation. It has 4 free parameters (as in **Eq. 1**) and is fit separately to the three different blocks (50:50, 80:20, 20:80), thus resulting in a total of 12 parameters. The *omniscient* and *fixed* models possess the perceptual decision-making model but do not compute the prior, *P*(*L*). Instead, the *omniscient* model has direct access to the true value (0.2, 0.5, or 0.8), and the *fixed* model uses a single *P*(*L*) throughout the duration of the session. Next, the *change-point* (CP) models (5 in total, see below) employ variants of the Bayes optimal online change-point detection model for estimating the prior, and the perceptual decision-making model for making a choice. The CP model has a potential of 4 free parameters. The a-priori block probabilities ξ*_t_* experimentally belong to the set *S*_1_ = {0.2, 0.5, 0.8}. In model fitting we set *S*_1_ = {*plow*, 0.5, *phigh*. Similarly, the run lengths are experimentally defined by a time constant τ = 60 and a minimum length *r*_NO7_ = 20 (**Eq. 15**). In the *change-point* model the 4 parameters (*plow*, *phigh*, τ, and *r*_NO7_) are set to the experimentally imposed values. In *CP free sym*, we set τ and *r*_NO7_ to their experimental values, and *plow* and *phigh* have to be symmetric (i.e., adding to 1). In *CP free* we allow *plow* and *phigh* to vary independently. *CP free sym run* is as *CP free sym* but also allows τ and *r*_NO7_ to take on any value. *CP free run* is as the *CP free* model but allows τ and *r*_NO7_ to take on any value. Finally, the exponential weighting models (2 in total, see below), use the exponential averaging for estimating the prior, and the same perceptual decision making model as the rest. In the no bias variant of the model (*exp. no bias*) the free parameter is α*_exp_*, dictating the shape of the time constant of memory decay. In the biased version of the model (*exp. bias*) there is additionally the parameters α and β dictating the shape of the Beta distribution acting as pseudo-counts.

We fit the above-described models by minimizing the negative log likelihood of the data using Bayesian Adaptive Direct Search (BADS^75^), and taking the best result of 20 optimization runs with randomized starting points. Further, we cross-validate log likelihoods (Fig. 2A) by splitting training and testing data according to odd and even sessions.

### Neural Analyses

#### Demixed PCA

We used demixed principal component analysis (dPCA^40^) to examine interpretable neural manifolds. This technique (see Kobak et al., 2016 for detail) requires convolved firing rates (as opposed to spike trains) and a given set of experimental conditions (i.e., not a continuous value). Thus, we convolved spike trains (1ms bins) with a causal Gaussian kernel (sd = 10ms), epoched each trial from 500ms prior to stimulus onset to 1500ms after stimulus onset, and averaged according to choice (left or right) and subjective prior. The latter, being a continuous variable, was binned in quintiles (5 levels of equal number of trials). The resulting matrix used in the dPCA was N (units) x P (5 quintiles of the prior) x D (2 choices) x T (time). While the analyses is conducted on individual CCF-defined regions, in the main text we amalgamate results across ‘macro-areas’ (**Table S1**) for statistical power and deriving a coarse-level summary. We included in the analysis sessions with at least 10 simultaneous units within a given CCF-defined region^39^. Including sessions/areas with more than 10 units simultaneously recorded results in a better estimate of the underlying latent dynamics but did not statistically change the fraction of variance explained by the different subspaces.

#### Poisson generalized additive model (pGAM)

To estimate tuning functions and their statistical contribution to a unit’s overall response, we fit a Poisson generalized additive model (P-GAM^42^). The P-GAM defines a non-linear mapping between spike counts of a unit *y_t_* ∈ ℕ_P_ and a set of continuous covariates *x*_j_, as well as discrete events *z*_k_. In this case, the continuous covariates included were both the experimentally imposed prior (e.g., 80:20 or 20:80 probability of a stimuli being on the left) and the “subjective” prior estimated via behavioral model fitting (exp. bias model). Further, to account for idiosyncratic body movements (Musall et al., 2019), we also included the first 10 principal components of video recordings. These PCs accounted for 79.16% of the variance in video data. As discrete events, we included visual contrasts ([−100%, −25%, −12.5%, −6.25%, 0%, 6.25%, 12.5%, 25%, and 100%]) at stimulus onset, as well choice and feedback (at their respective times), both for the current trial as well as for the previous one. The previous choice and feedback were modeled as accounting for sustained responses (as opposed to evoked) even prior to trial onset. Lastly, above and beyond the experimental variables, we also accounted for elements of internal neural dynamics by including the concurrent firing of simultaneously recorded units in the same region, *y_t_*. Together, a unit’s log-firing rate is modeled as a linear combination of arbitrary non-linear functions (B-splines) of the covariates,

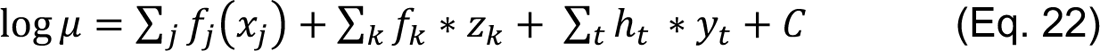

where * is the convolution operator, and the spike counts are generated as Poisson random variables with the rate specified by (**Eq. 22**). Input specific non-linearities *f*(⋅) were expressed in terms of flexible B-splines, *f*(⋅) ≈ β ⋅ *b*(⋅). Similarly, ℎ_(_ are smooth causal filters (also learned) capturing the directional coupling between units, including an auto-regressive component which accounts for refractory periods of units. Covariates and spike counts were discretized in 5ms bins. The estimated kernels (*f* and ℎ) were associated with a smoothness enforcing penalization term controlled by a scale parameter λ_S_,

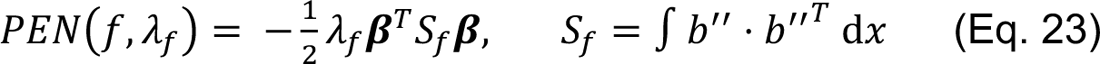

The larger λ_S_, the smoother the model. These penalization terms can be interpreted as Gaussian priors over model parameters. The resulting log-likelihood of the model takes the form,

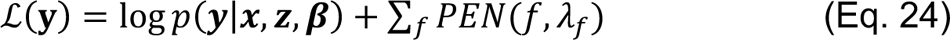

with *y* ∈ ℝ^*T*^ being the spike counts of the unit, *x* ∈ ℝ^V×T^ being the continuous task variables, *z* ∈ ℝ^X×T^ being the discrete task events, *T* being the time points, β being the collection of all B-spline coefficients, and being *p*(⋅) the Poisson likelihood. Both parameters β and the hyperparameters λ are learned from the data by an iterative optimization procedure that switches between maximizing **Eq. 24**, and minimizing a cross-validation score as a function of hyperparameters (see^42^, for further details). We used 11 nodes (βs) per fitted co-variable (19 experimental + a variable number of simultaneously recorded units), resulting in the average full encoding model having 590.33 parameters. Most importantly, the probabilistic interpretation of the penalization terms allowed us to compute a posterior distribution for the model parameters. In turn, this allows us to derive confidence intervals with desirable frequentist coverage properties and implement a statistical test for inclusion of a minimal subset of task variables explaining most of the variance. In other words, it allows fitting an encoding model and performing implicit model selection within our large neurophysiological dataset (see Noel et al., 2022, 2023 for a similar approach and additional model validations). The average reduced model had 61.25 parameters (10.3% of the full model) with no detriment to its ability to account for spiking activity (all p > 0.36, **Fig. S6**).

The encoding model fit quality was assessed via pseudo-R^2^ on subset of held-out test trials (20% of the total trials). Pseudo-R^2^ is a goodness-of-fit measure that is suitable for models with Poisson observation noise^76^. The score is computed as,

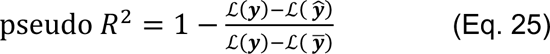

with ℒ(*y*) being the likelihood of the true spike counts, ℒ( *y*Å) being the likelihood of the pGAM prediction, and ℒ( *y*Ç) being the likelihood of a Poisson null-model (mean rate). Pseudo-R^2^ is 0 when the pGAM fits are no more likely than the null model, 1 when it perfectly matches the data, and can be negative when overfitting occurs (for test-set data, 0.5% of the recorded units). Empirically, the pseudo-R^2^ is a stringent metric and ranges in values that are substantially lower than the standard R^2^ when both are applicable^77^. Our average score of 0.0745 is two to four times better than standard GLM performance^47^. The R^2^ of trial-averaged firing rates to variables deemed to significantly contribute to spike trains was on average 0.82. For analyses downstream of the pGAM, we include units with a minimum pseudo-R^2^ of 0.01, deem variables as significantly contributing to spike trains at alpha < 0.001, and include regions in the analysis if at least 40 units per area were properly fit, in each of the genotypes (Fig. 5, Fig. 6C**-E**) or across genotypes (Fig. 6A, B). For Figure 7, we fit separate pGAMs for each biased block (80:20 and 20:80) and remove the experimental prior as a co-variate. This allowed us to estimate responses to each contrast for each of the experimental priors.

We compute the mutual information between an experimental variable and the observed spiking activity, where *y_t_* are the spike counts at time *t*, and *y_t_*|β, *X_t_* ∼ *Poisson*(*X_t_*, β) where *X*_(_ is the stimulus matrix at time *t*. We know that β| *X* ∼ *N*(β^Ö^, Σ). We select one stimulus dimension, *x*_Q_ = *X*_:Q_, and discretize it into N values, *x*_Q_ ∈ {*x*^&^ … *x*^Y^}. We approximate the input distribution using a binomial *p*(*x*^R^) = *p*_R_, where *p*_R_ is the empirically observed frequency of the stimulus. For each time point we computed,

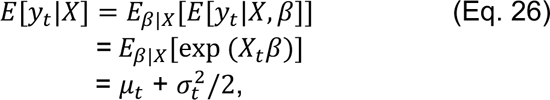

Where μ*_t_* = *X_t_*β^Ö^ and σ^2^=*X_t_*Σ *X*^T^. We approximate the conditional entropy *H*[*y*|*x*^R^] as the entropy of a Poisson variable with mean equal to & Σ*_t_H*[*y_t_*|*X*], where the sum is taken over the T time points in which *x*_Q*t*_ = *x*^R^. We also approximate the entropy of spikes *H*[*y*] as the entropy of a Poisson with mean the average firing rate of the unit. Mutual information was computed as,

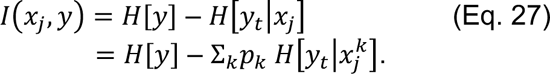

## Supplementary Materials

**Figure S1.**
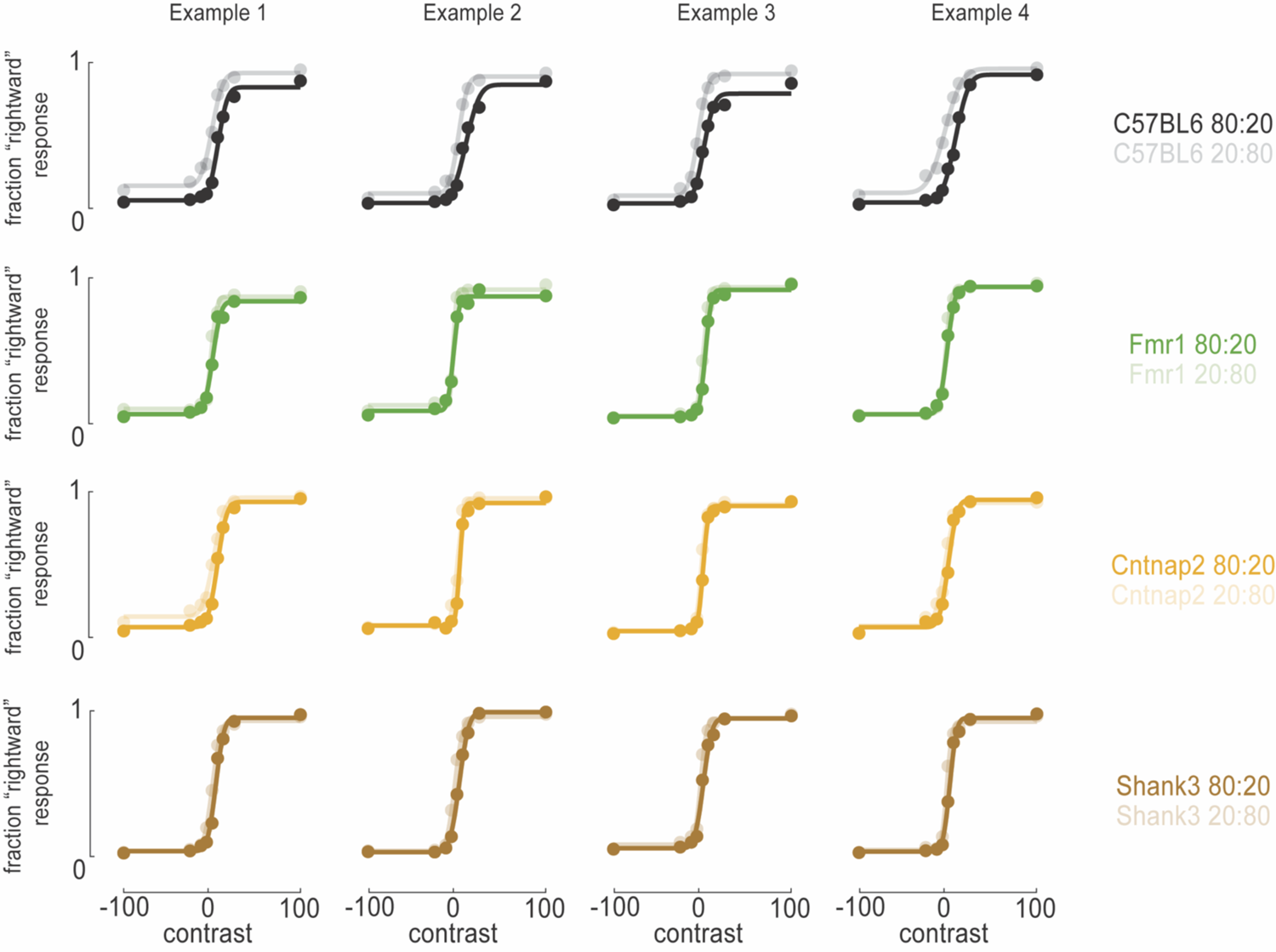
Example psychometric fits during biased sessions. Psychometric fits to the fraction of “rightward” responses as a function of contrast (x-axis) and experimental block (colors; dark color indicated a left-biased block). Four example animals (columns) are shown for each of the 4 genotypes (rows). Circles are the observed fraction of responses, and curves are fits.

**Figure S2.**
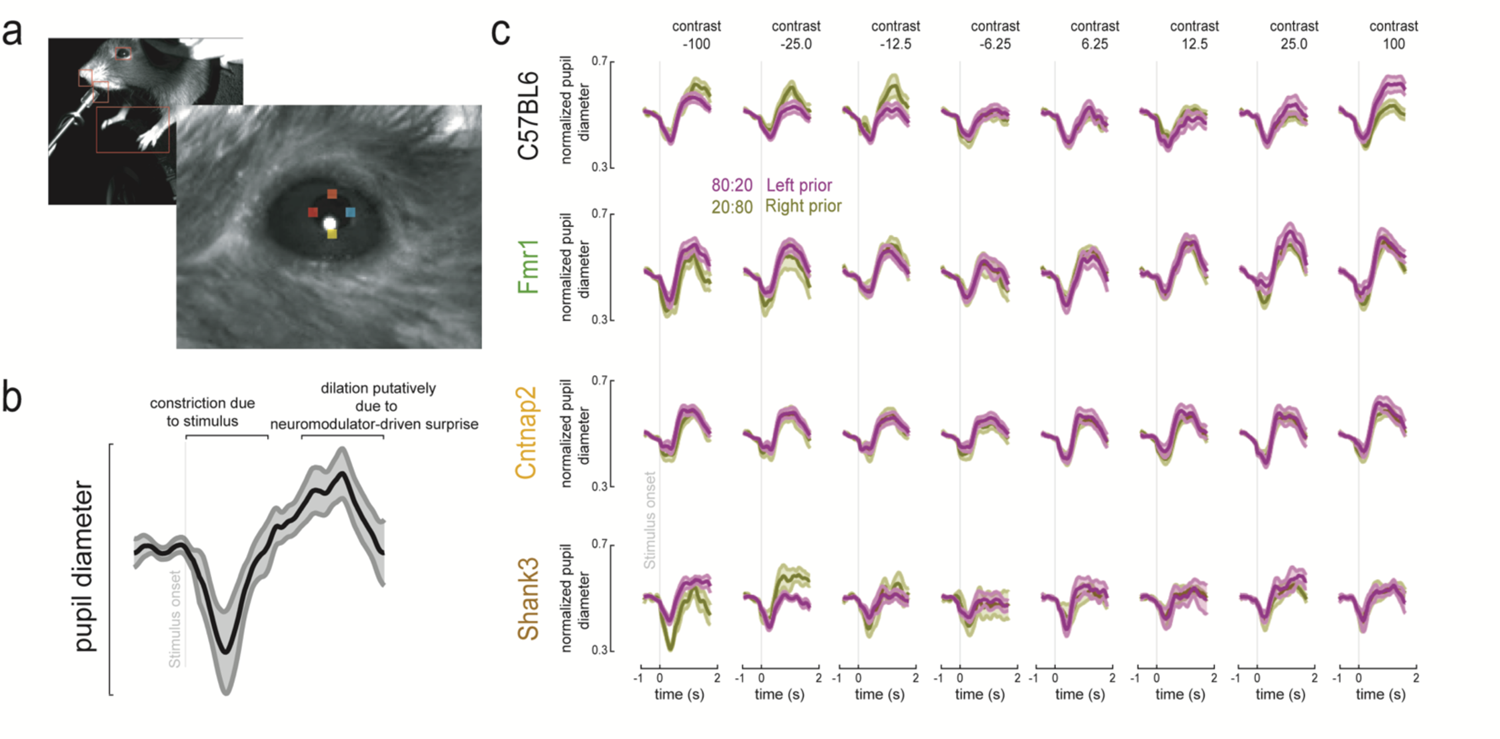
Lack of surprise during the presentation of statistically unlikely events in mouse models of ASD. **A.** Video recordings and schematic showing pupil tracking. **B.** Grand-average pupil diameter as a function of time since stimulus presentation. **C.** Pupil diameter as a function of experimental block (80:20 in purple, 20:80 in gold), genotype (rows; C57BL6, Fmr1, Cntnap2, and Shank3, respectively) and contrasts (columns). In control animals (C57BL6) we observe that at −100% (p < 0.05, 1.79s post-stimulus onset), −25% (1.31s post-stimulus onset), and 100% (1.23s post-stimulus onset) contrasts, the late latency (i.e., surprise-driven) pupil diameter was modulated by sensory history. Importantly, dilation was greater when statistically unlikely events were presented (i.e., high contrast on the right visual field under the left prior, or high contrast on the left visual field during the right prior) and did not occur when sensory observations were uncertain (low contrast). The prior-dependent pupil dilation indicating surprise was not present in any of the mouse models of ASD (second and third row, all p > 0.16), with exception for a single contrast (−25%, p < 0.05, 1.46 post-stimulus onset) in the Shank3 animals (bottom row, second column). Shaded area is ± 1 s.e.m.

**Figure S3.**
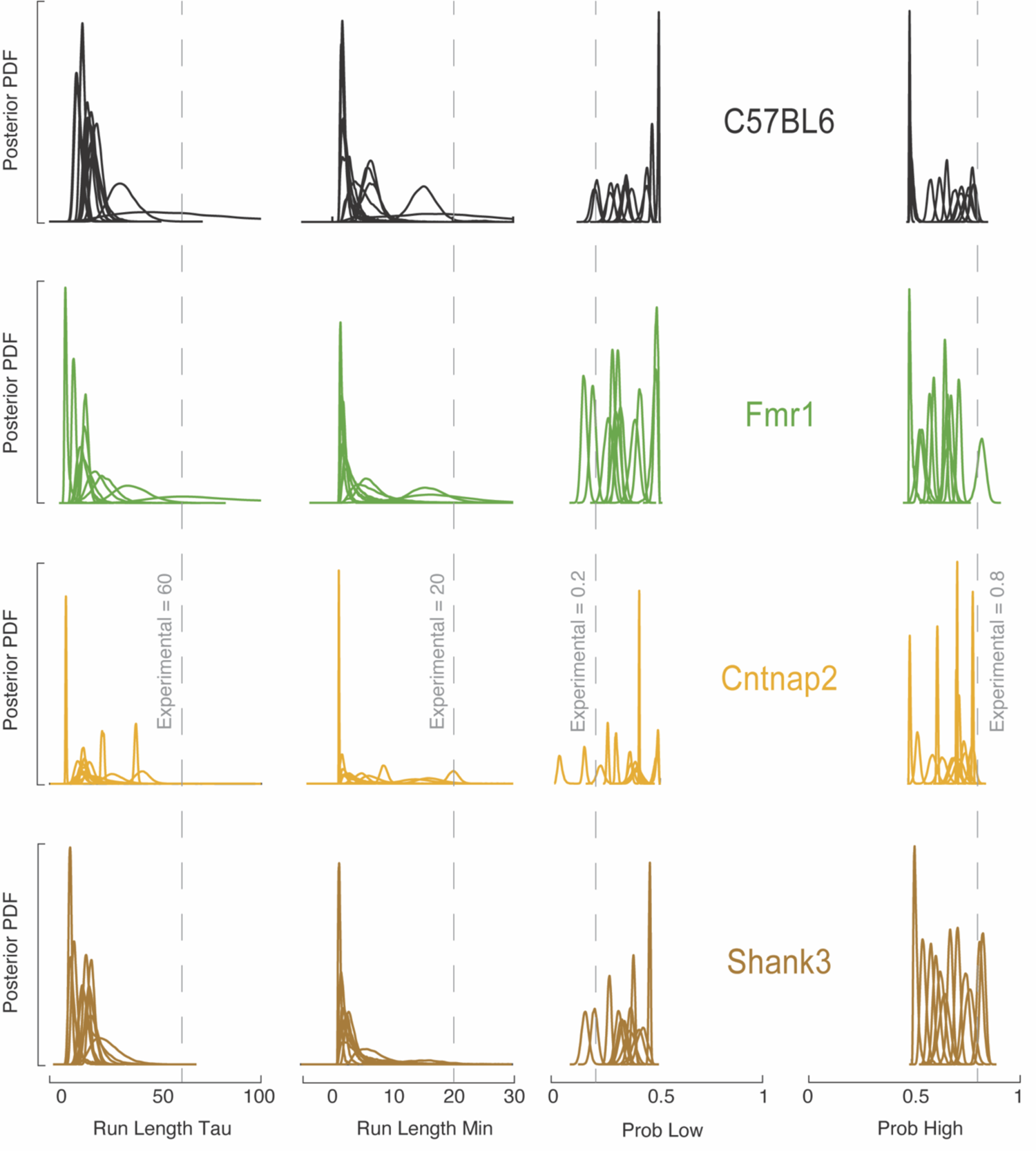
We fit the Bayesian *change-point free run* model – the most flexible of the change-point models – to the animals behavioral responses. This model performs worse than the biased weighted average in general (i.e., when considering all animals), and statistically for the Cntnap2 and Shank3 animals. It performs equally to the biased weighted average model in C57BL6 and Fmr1 animals. Here, we estimate full posteriors of the *c.p. free run* model via Variational Bayesian Monte Carlo (VBMC^78^). These, demonstrate that the posteriors were generally well-behaved (i.e., sufficiently narrow not to encompass the full parameter-space), yet far from the true experimental values. In particular, it is worth noting that the time constant and minimum run lengths estimated by animals were very concentrated near 0. In other words, animals did not use a generative model wherein there was a representation of blocks. Instead, the model parameters attempted to exclude this notion by making “blocks” as short as possible, effectively rendering this *c.p. free run* model akin to the heuristic models detailed in the main text.

**Figure S4.**
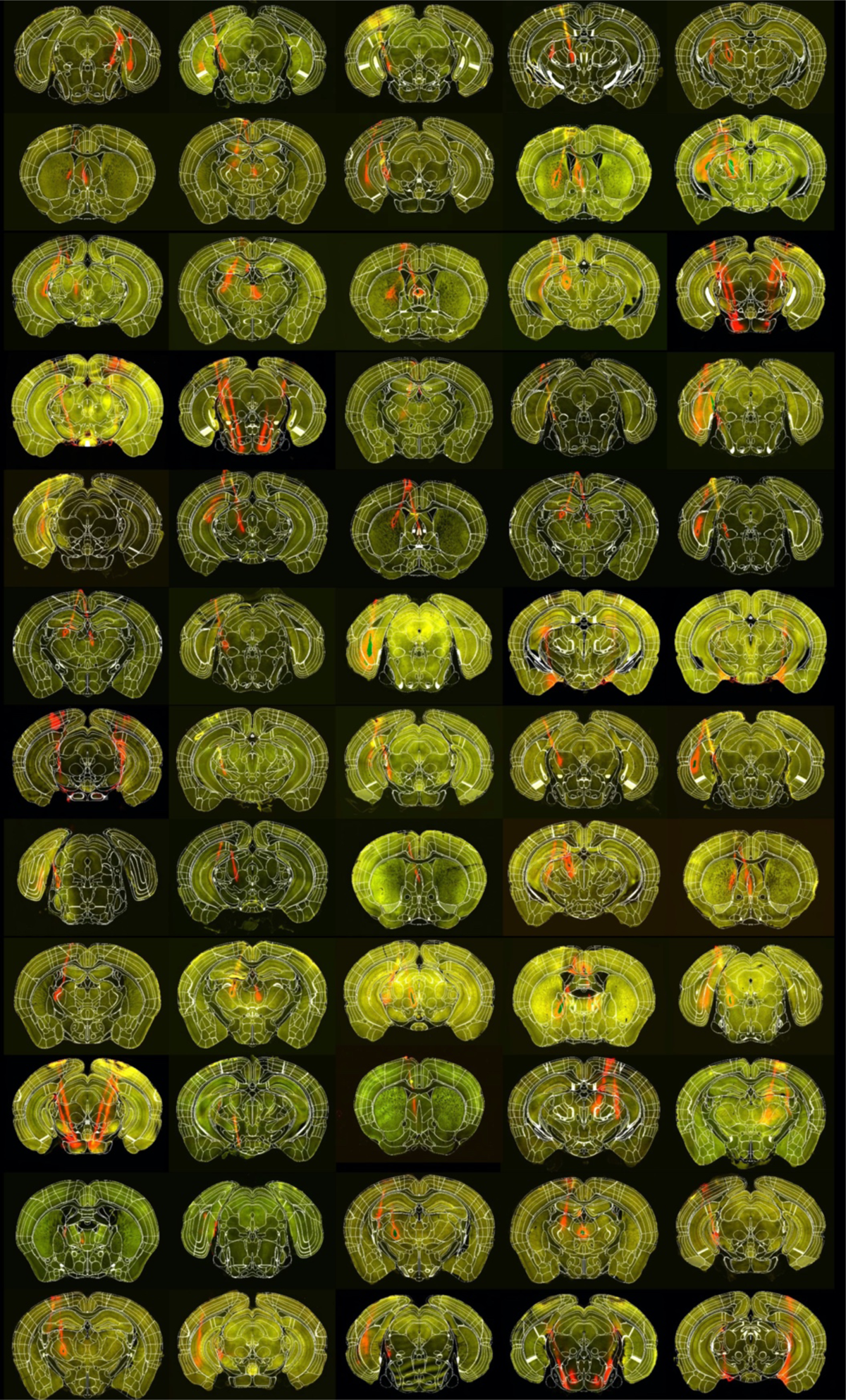
Electrode track histology with CCF-atlas alignment overlaid. Sixty coronal histology slices are shown. The dye is shown in red, tracking the electrode tracks. Allen Common Coordinate Framework (CCF) atlas is shown overlayed in white. Tracing these tracks rendered the problem of localizing units from 3-dimensional (in the brain), to 1-dimensional (along the track). Histology and physiology were then aligned (Falkner, 2020) the track to solve the 1-dimensional problem.

**Figure S5.**
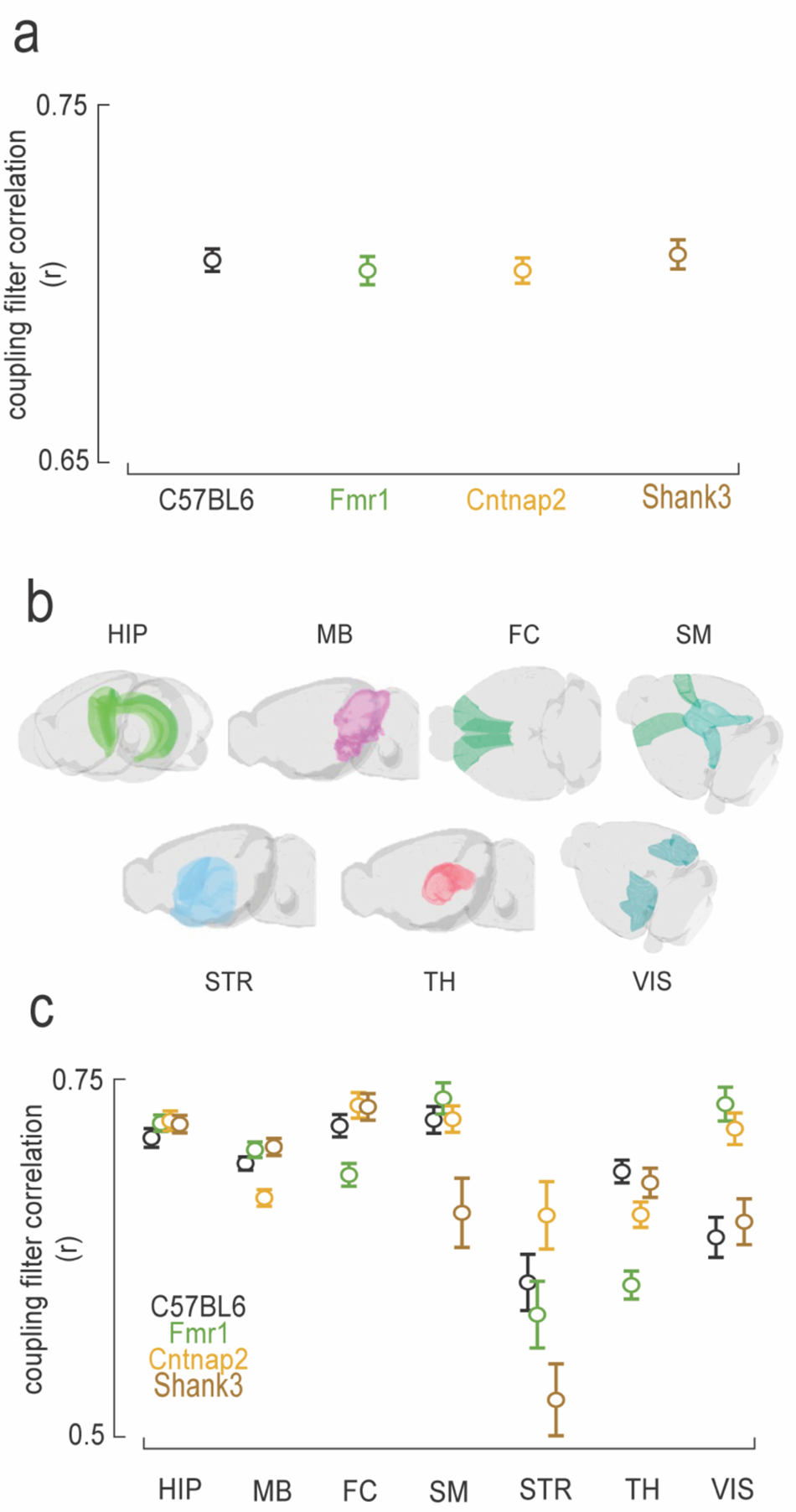
Stability of coupling filters across experimental blocks. **A.** We compute the Pearson correlation coefficient (r) between coupling filters across a given pair of units estimated by the pGAM in leftward (80:20) and rightward-biased (20:80) experimental blocks (over 640k pairs in total). These showed a strong degree of stability (r ∼ .70) and no difference across genotypes (One-way ANOVA, p = 0.79). When separating into macro-areas (panel **B**), we observed more remapping of noise correlations in STR than the rest of areas (p < 0.05, panel **C**), but no systematic effect wherein all mouse models of ASD differed from the control.

**Figure S6.**
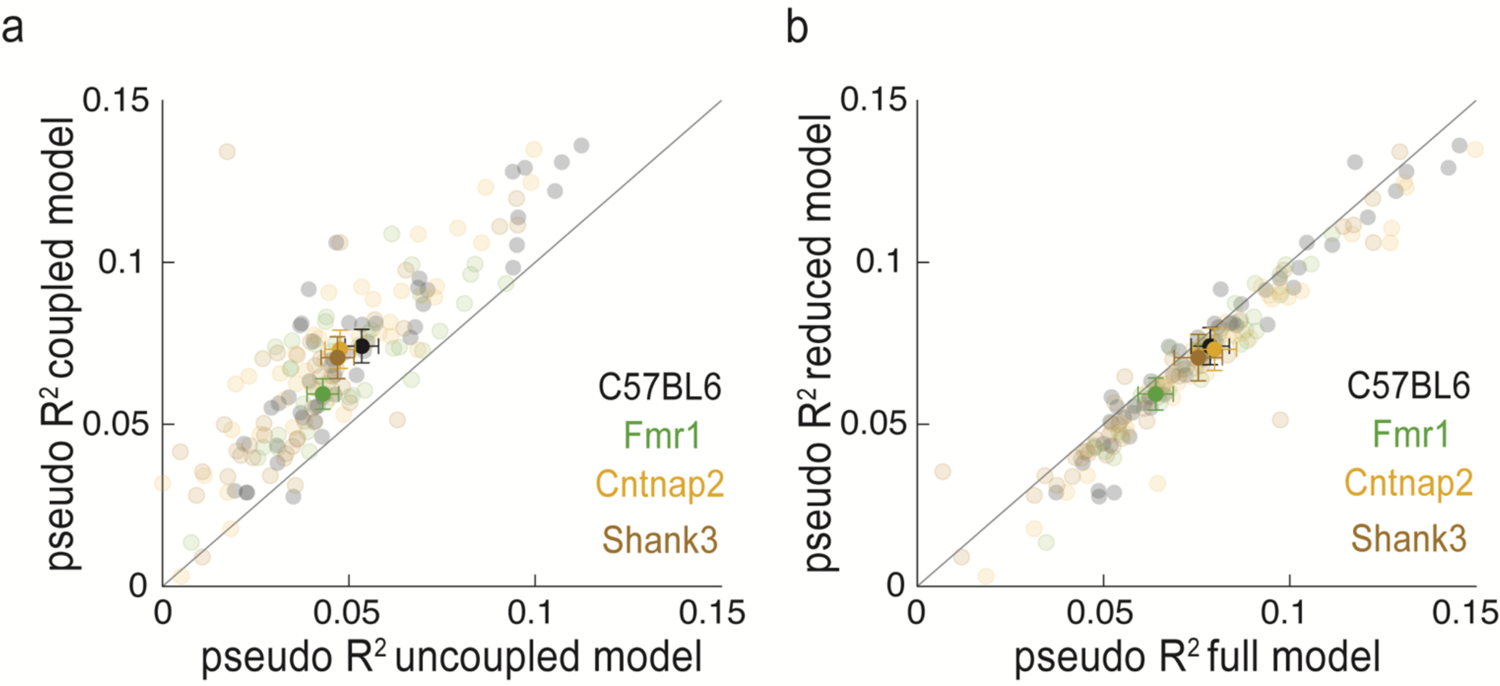
Performance of the encoding model (pGAM). We fit a coupled model (i.e., with neural responses of one unit putatively impacting the firing of another) in order to account not solely for task driven responses, but also for internal neural dynamics. Left panel: comparison of pseudo-R^2^ for the coupled model (y-axis) and an uncoupled model (x-axis). Transparent dots are sessions (colored according to genotype), and opaque dots are averaged across sessions. Error bars are S.E.M. In all genotypes, the coupled model accounts better for spike trains (all p < 0.003). To assess performance of the variable selection procedure, we contrast pseudo-R^2^ of the models allowing for coupling, in full (x-axis, average of 590.33 parameters) and when reduced to the variables deemed to significantly accounting for spiking activity (y-axis, average of 61.25 parameters, alpha set at 0.001). The full and reduced model accounted for an equal portion of the variance (all p > 0.36), while the latter had a tenth the number of parameters/retained variables.

**Figure S7.**
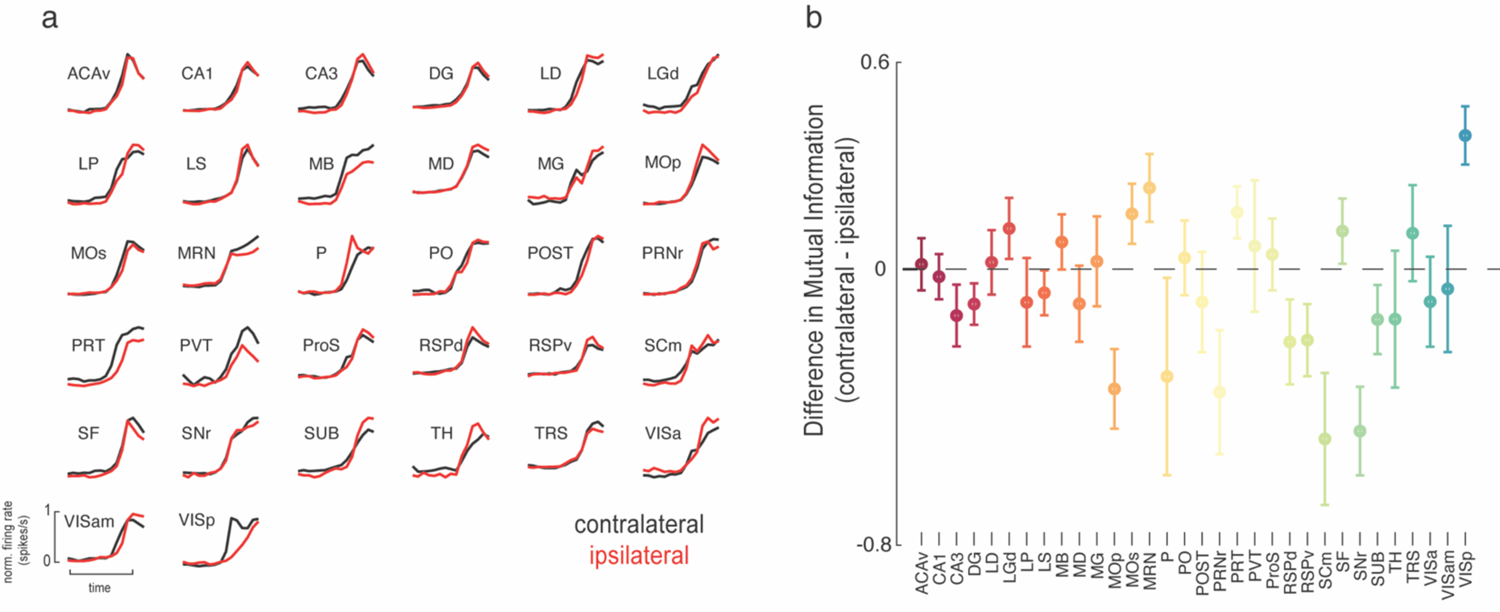
Responses to contra- and ipsi-lateral high contrast gratings predicted by the pGAM. **A.** Evoked firing rates to high contrast gratings in contra-(left, black) and ipsi-(right, red) lateral visual field as a function of brain region. As previously demonstrated (Steinmetz et al., 2019) primary visual cortex (VISp) is particularly tuned to contra-lateral stimuli, while the rest of regions respond fairly equally across hemi-fields. **B.** Difference in the mutual information between neural responses evoked by contra- and ipsi-lateral grating presentation.

**Figure S8.**
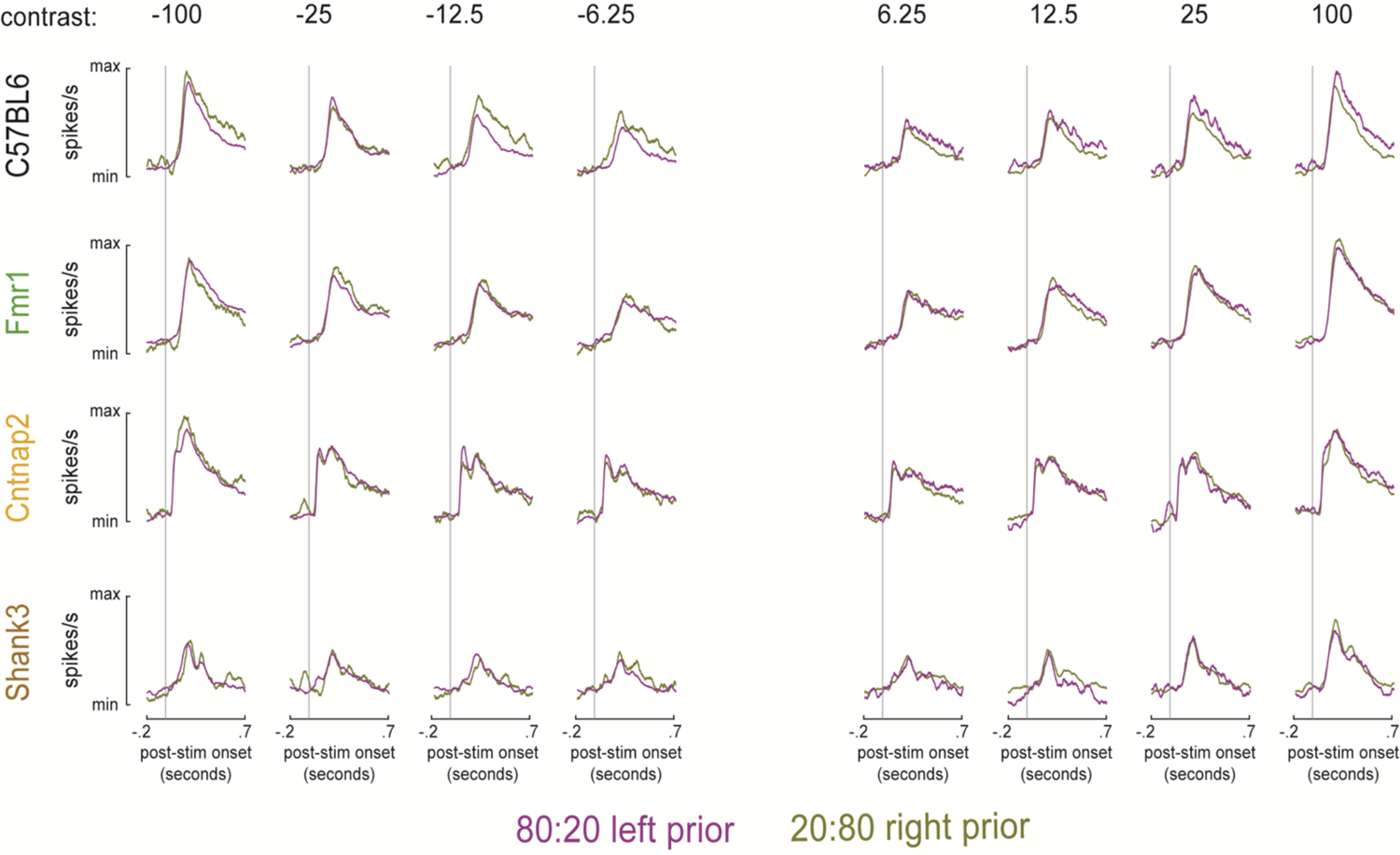
Peri-Stimulus Time Histograms (PSTH) across frontal cortices as a function of sensory history. In control animals (C57BL7, top row) neural responses were stronger to unexpected stimuli – for instance, a grating on the left hemifield (negative contrasts) under a rightward-biased block (gold color). This effect, consistent with predictive coding, was absent in Fmr1 (second row, green), Cntnap2 (third row, yellow), and Shank3 (brown) animals.

---

**Table S1.**
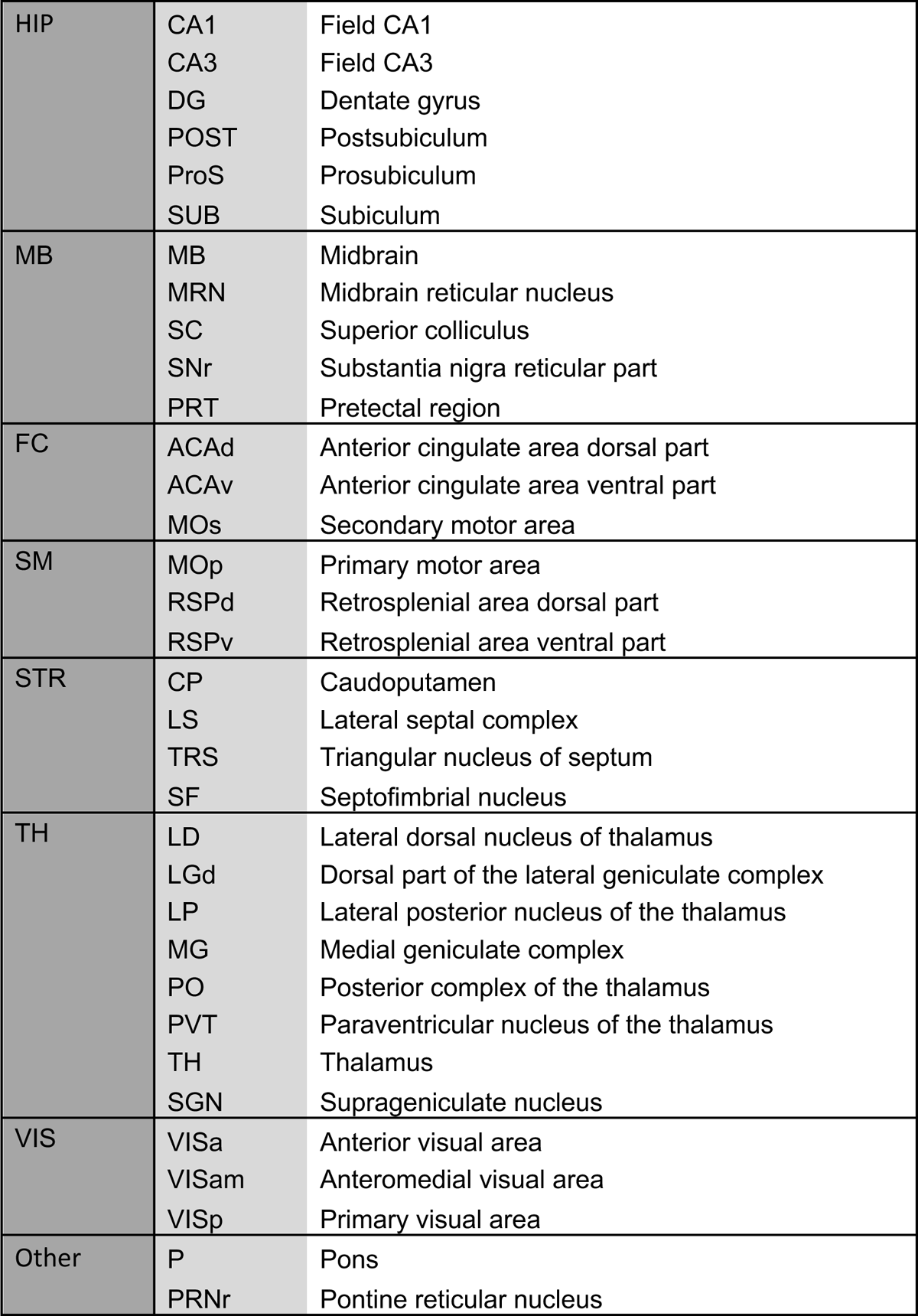
Categorization of areas and their acronyms.

